# LncRNA Snhg1-driven self-reinforcing regulatory network promotes cardiomyocyte cytokinesis and improves cardiac function

**DOI:** 10.1101/2020.10.26.354613

**Authors:** Mengsha Li, Hao zheng, Yijin Chen, Bing Li, Guojun Chen, Xiaoqiang Chen, Senlin Huang, Xiang He, Guoquan Wei, Tong Xu, Xiaofei Feng, Wangjun Liao, Yulin Liao, Yanmei Chen, Jianping Bin

## Abstract

Most of current cardiac regenerative approaches result in very limited cell division. Positive feedback loops are vital for cell division, but their role in CM regeneration remains unclear. We aimed to demonstrate that lncRNA Snhg1 formed a positive feedback loop with c-Myc to induce stable CM cytokinesis. We found that Snhg1 expression was increased in human and mouse fetal and myocardial infarction (MI) hearts, particularly in CMs. Snhg1 overexpression elicited stable CM proliferation and improved post-MI cardiac function. Antagonism of Snhg1 in neonatal mice inhibited CM proliferation and impaired cardiac repair after MI. Proliferative effect was confirmed using cardiac-specific transgenic mice. RNA pull-down assays showed that Snhg1 directly bound to PTEN and activated PI3K-Akt pathway, resulting in c-Myc activation. Chromatin immunoprecipitation experiments showed that Snhg1 expression was upregulated by c-Myc binding to the Snhg1 promoter region, indicating a positive feedback loop between c-Myc and Snhg1. In conclusion, c-Myc/Snhg1/PI3k-Akt positive feedback loop drove sustained activation of cell cycle re-entry and induced stable CM cytokinesis, and thus may be an attractive strategy for promoting heart regenerative response.

**Clinical Perspectives:** Most of the current cardiac regenerative approaches result in very limited cell division and little new cardiomyocyte (CM) mass. Positive feedback loops are vital for cell division, but their role in CM regeneration remain unclear. Here, we identified the long noncoding RNA Snhg1 as a driver to induce stable CM division and improve cardiac function after myocardial infarction (MI) by forming a positive feedback loop to sustain PI3K-Akt signaling activation. This finding might provide a novel therapeutic of Snhg1 as a promising regenerative approach to improve the prognosis of patients with heart failure.

## Introduction

Adult mammalian cardiomyocytes have limited regenerative capacity to replace the lost cardiac tissue after acute ischaemic injury (Eschenhagen *et al*, 2017). The mammalian cardiomyocytes withdrawal from cell cycle shortly after birth and are incapable of undergoing cell division (cytokinesis), resulting in binucleation (Derks & Bergmann, 2020; Karra & Poss, 2017). Hence, a scientific and clinical imperative is to find regenerative strategies that direct CMs to undergo efficient cytokinesis to form a new daughter cell. Successful cytokinesis requires efficient progression through both G1/S phase and G2/M phase (Fededa & Gerlich, 2012). A recent study established a novel combination of cyclin-dependent kinase 1 (CDK1), CDK4, cyclin B1, and cyclin D1 which efficiently induced cell division in post-mitotic cardiomyocytes (Mohamed *et al*, 2018). To control cell division, multiple positive- and/or negative-feedback loops are also indispensable (Santos *et al*, 2012). Prolonged exposure to cell cycle regulators is required for cell cycle entry and commitment to cell division (Santos *et al.*, 2012; Yoshizumi *et al*, 1995). A positive feedback loop is a self-reinforcing loop that sustains the activation of cell cycle re-entry and contributes to the irreversibility of cell division (Skotheim *et al*, 2008). A recent study showed that a c-Myc-driven positive feedback loop triggered epigenetic memory in embryonic stem cells to maintain their self-renewal capacity (Fagnocchi *et al*, 2016). However, the roles of the positive feedback loop in the induction of CM cytokinesis and heart regeneration remain unknown.

Long noncoding RNAs (lncRNAs) are a class of RNA transcripts over 200 nucleotides in length and without protein-coding potential. Previous studies and our studies have shown that several lncRNAs can promote CM proliferation (Cai *et al*, 2018; Chen *et al*, 2019). The lncRNA small nucleolar RNA host gene1 (Snhg1; Gene Bank: 23642 (human); 83673 (mouse)), which was originally identified by its oncogenic activity (Thin *et al*, 2019), was shown to promote cancer cell proliferation by activating the PI3K/Akt signaling pathway (Sun *et al*, 2017; Zhang *et al*, 2018). PI3K/Akt signaling has also been implicated in the control of heart regeneration (Engel *et al*, 2005; Gabriele *et al*, 2015). Therapeutic targeting of upstream signaling pathways that regulate cell proliferation may reduce the risk of teratogenicity (Bersell *et al*, 2009). We predicted that Snhg1 can function upstream of PI3K/Akt signaling to control CM cytokinesis and trigger heart regeneration. Activation of PI3K/Akt signaling induces the expression of CDK regulatory proteins, including c-Myc (Amit & Ana C, 2007), and online tools were used to predict that c-Myc binds to the Snhg1 promoter. Thus, there may be a positive feedback loop between c-Myc and Snhg1.

In this study, we predicted that c-Myc and Snhg1 form a positive feedback loop to sustain activation of PI3K/Akt signaling, which may induce CM cytokinesis. Once established, this Snhg1-driven self-reinforcing circuit may direct CMs to undergo cytokinesis as opposed to karyokinesis, representing a powerful approach for mammalian heart regeneration.

## Methods

### Animal model

Healthy C57BL/6J mice were purchased from the Laboratory Animal Center of Southern Medical University (Guangzhou, China). The Cre-dependent Cas9 knock-in mouse model (R26-CAG-Cas9/+) was purchased from Shanghai Model Organisms Center, Inc. The Myh6-mCherry transgenic mice (Myh6*-mCherry R26-mTmG) were kindly provided by Dr. Jinzhu Duan from Guangzhou Women and Children’s Medical Center, China. All animal procedures were conformed to the guidelines from Directive 2010/63/EU of the European Parliament on the protection of animals used for scientific purposes or the NIH Guide for the Care. Approval for this study was granted by Southern Medical University’s ethics review board.

### Establishment of MI or myocardial ischaemia/reperfusion (I/R) model

Mouse MI was carried out as described previously.(Chen *et al.*, 2019) P1 and P7 mice were anesthetized by cooling on an ice bed for 6 min, whereas adult male mice (8-10 weeks of ages) were intraperitoneally anesthetized with 3% pentobarbital sodium (40 mg/kg) following tracheal intubation for artificial ventilation. An ALC-V8S rodent ventilator (ALCBIO, Shanghai, China) was used to supply oxygen during the surgical procedure. The chests were opened by a horizontal incision through the muscle between the fourth and fifth intercostal space. The pericardium was then removed. The LAD coronary artery was permanently ligated with a 9-0 silk suture (Ningbo Medical Needle Co., Ningbo, China). After surgery, the thoracic wall and skin were closed with a 5-0 silk suture. Myocardial ischemia was confirmed by an electrocardiogram ST-segment elevation using an Animal Bio Amp (Ad instruments, NSW, Australia). After ligation, AAV9 vectors were injected immediately into the myocardium bordering the infarct zone using an insulin syringe with a 30-gauge needle. After surgery, the skin was disinfected, and the animals were revived while being kept on a thermal insulation blanket. The hearts were collected at 14 or 28 days after infarction as described below.

For the model of myocardial I/R, after occlusion of the left coronary artery for 45 min, the ligature was released for myocardium reperfusion. Successful reperfusion was confirmed by observing the return of a bright red color to the pale region in the myocardium. Sham control mice underwent the same anesthetic and surgical procedures, except the ligation of the LAD was not tied.

### Tissue collection

Mice were anesthetized with 2% isoflurane and then sacrificed by injection of 10% KCl. The hearts and lungs were excised, briefly washed with 0.9% NaCl, weighed, and fixed in 10% formalin at room temperature. The hearts were embedded in paraffin and further processed for histology or immunofluorescence analysis.

### 1-day-old and 7-day-old mouse ventricular CM isolation

1-day-old C57BL/6J mice were anesthetized with 2% isoflurane inhalation and sacrificed by cervical dislocation. The ventricles from neonatal mice were separated from the atria, cut into pieces, and digested with 0.25% trypsin (Gibco, Grand Island, NY, USA) at 4°C overnight. Digestion was repeated in collagenase type II (Gibco) with bovine serum albumin (BSA) (Sigma, St. Louis, MO, USA) in phosphate-buffered saline (PBS) at 37°Cfor 15 min twice under constant stirring. Digestion was performed at 37°C in 15-min steps, and the supernatant was collected and mixed with fetal bovine serum (FBS) (Gibco) after each step. The collected supernatant was centrifuged to separate the cells, which were resuspended in Dulbecco’s modified Eagle’s medium/nutrient F-12 Ham (DMEM/F12) 1:1 medium (HyClone, Logan, UT, USA) supplemented with 10% FBS and 100 U/mL penicillin (Sigma) and 100 mg/mL streptomycin (Sigma). The collected cells were seeded onto 100-mm plastic dishes for 2h at 37°C in a humidified atmosphere of 5% CO2. The supernatant, composed mostly of CMs, was collected and pelleted. Cells were resuspended in DMEM/F12 containing 10% FBS media, penicillin, and streptomycin, and then counted and plated at an appropriate density.

Ventricular CMs from 7-day-old mice were isolated essentially as described for neonatal mice, except that the digestions were performed at 37°C in calcium- and bicarbonate-free Hanks’ solution with HEPES (CBFHH) buffer containing 0.25 mg/ml of pancreatin (Sigma), 0.125 mg/ml collagenase type II (Worthington Biochemical) and 10 mg/ml DNase II (Sigma), under constant stirring. Cells were washed thoroughly 24h after seeding, and transfection.3

### Adult mouse CM isolation

Adult CMs were isolated from 6-to 8-week-old mice as previously described.(Huang *et al*, 2019) Briefly, adult mouse hearts were anesthetized with 2% isoflurane and their hearts were dissected and perfused using solution A (118 mM NaCl, 4.8 mM KCl, 25 mM HEPES, 1.25 mM MgSO4, 1.25 mM K2HPO4, 10 mM glucose, 4.95 mM taurine, 9.89 mM 2,3-butanedione monoxime; pH 7.35). The hearts were then adjusted on a Langendorff perfusion system and digested with digestion buffer (solution A with 0.1% BSA, 0.05 mM CaCl2, 0.07% collagenase type II, 0.02% hyaluronidase type I). Next, ventricular tissue was removed and minced in digestion buffer. The cell suspension was filtered through a 100-μm cell strainer (BD Biosciences, Franklin Lakes, NJ, USA), and the filtrate was centrifuged for 3 min at room temperature. The cell pellet was resuspended in solution B (solution A with 1% BSA and 0.1 mM CaCl2) and the cells were allowed to settle under gravity. This cell pellet was resuspended and seeded into DMEM/high glucose media (HyClone) containing 10% FBS, 100 U/mL penicillin, and 100 mg/mL streptomycin.

### Adenovirus (Adv) or siRNA transfection in vitro

Adv-containing green fluorescent protein (GFP) vectors for Snhg1 overexpression and si-Snhg1 were synthesized by Vigene (Shandong, China). The sequence of Snhg1 siRNA was as follows: 5’-GCATTCAAAGGTTCTGTTATT-3’; 5’-TAACAGAACCTTTGAATGCTT-3’. The siRNAs of c-Myc and Gsk3β were synthesized by RiboBio (Guangzhou, China). The target sequence of c-Myc siRNA was: 5’-CTATGACCTCGACTACGAC-3’. The target sequence of Gsk3β siRNA was:5’-GUCCUAGGAACACCAACAA-3’. The PTEN inhibitors LY294002 were purchased from Calbiochem (Merck Eurolab, Fontenay Sous Bois, France).

Isolated mouse CMs were seeded at 70% confluence. Transfection was performed after 48 h of culture. Various MOI of adenovirus was added to cells. For siRNA transfection, 5μL Lipofectamine 2000 (Invitrogen, Carlsbad, CA, USA) and 50 nM siRNAs were added to Opti-MEM medium (Gibco). The mixed solution was incubated at room temperature for 20 min and then added to the cells. After 48 h, the cells were subjected to RNA or protein isolation or immunofluorescence analysis. The transfection efficiency of adenovirus and siRNA was determined by staining with an anti-GFP antibody (Biosynthesis, Beijing, China).

### Injection of AAV9 vectors in 7-day-old mice or adult mice

The AAV9-GFP vector overexpressing Snhg1 or NC was synthesized by Vigene. AAV9-mediated Snhg1 or NC were delivered by intracardiac injection into the left ventricle in 7-day-old mice at a dose of 2 × 10^10^ viral genome particles per animal using an insulin syringe with a 30-gauge needle (BD Biosciences). And the adult mice received the injection at a dose of 1 × 10^11^ viral genome particles per animal. The transfection efficiency of AAV9 was determined by staining with an anti-GFP antibody (Abcam, ab13970). In addition, In vivo imaging system (In-Vivo FX PRO; BRUKER Corporation, Billerica, MA, USA) was used to evaluate GFP fluorescence in 28 days after AAV9 injection. Adult mice were anesthetized and placed in the shooting position of In vivo imaging system. The fluorescence in vivo imaging system equipped with a LED that emitted light at a 480nm and a charge-coupled device (CCD) as an image detector was used to evaluate GFP fluorescence. In situ hybridization (ISH) and real-time polymerase chain reaction (RT-qPCR) assays were used to detect Snhg1 expression after transfection.

### Statistical analysis

All data are summarized as the mean ± standard deviation (SD). Statistical analyses were performed using SPSS 20.0 software (SPSS, Inc., Chicago, IL, USA). All data were subject to tests for normality. Data that do not follow a normal distribution were analyzed via a non-parametric equivalent. To statistically compare two groups, the unpaired, two-tailed Student’s *t*-test was used; to compare three or more groups, one-way or two-way analysis of variance (ANOVA) followed by the least significant difference (LSD) post hoc test was used. a two tail *P* <0.05 was considered as significant. Full methods are provided in the Supplemental Materials.

## Results

### Snhg1 is highly expressed in fetal and MI hearts

As the embryonic phase (measured at embryonic day 16.5 (E16.5)) progressed to adulthood (measured at postnatal day 56 (P56)), Snhg1 markedly decreased during heart development (Fig.1A) and showed most obvious change among Snhg family members (Fig.S1A). Snhg1 was conserved across humans, mice and rats (Fig.S1B), and was predominantly expressed in the liver, muscle, kidney and heart tissues (Fig.S1C). *In situ* hybridization (ISH) analysis showed that Snhg1 is highly expressed in both human and murine fetal hearts (Fig.B). The Snhg1 level in isolated neonatal mouse CMs was higher than that in cardiac fibroblasts (CFs) (Fig.1C). Costaining analysis both *in vitro* and *in vivo* revealed that Snhg1 expression in neonatal CMs was significantly higher than that in non-CMs (Fig.1D-E). After MI in adult hearts, Snhg1 expression increased in border and infarcted zone (Fig.1F-G), and this increase was mainly found in CMs but not in CFs (Fig.1H). Snhg1 was mainly located in the cytoplasm of neonatal mouse CMs but not in the nucleus (Fig.1I-J). Overall, these data demonstrated that Snhg1 was highly expressed in fetal and MI hearts, particularly in CMs.

**Figure 1.**
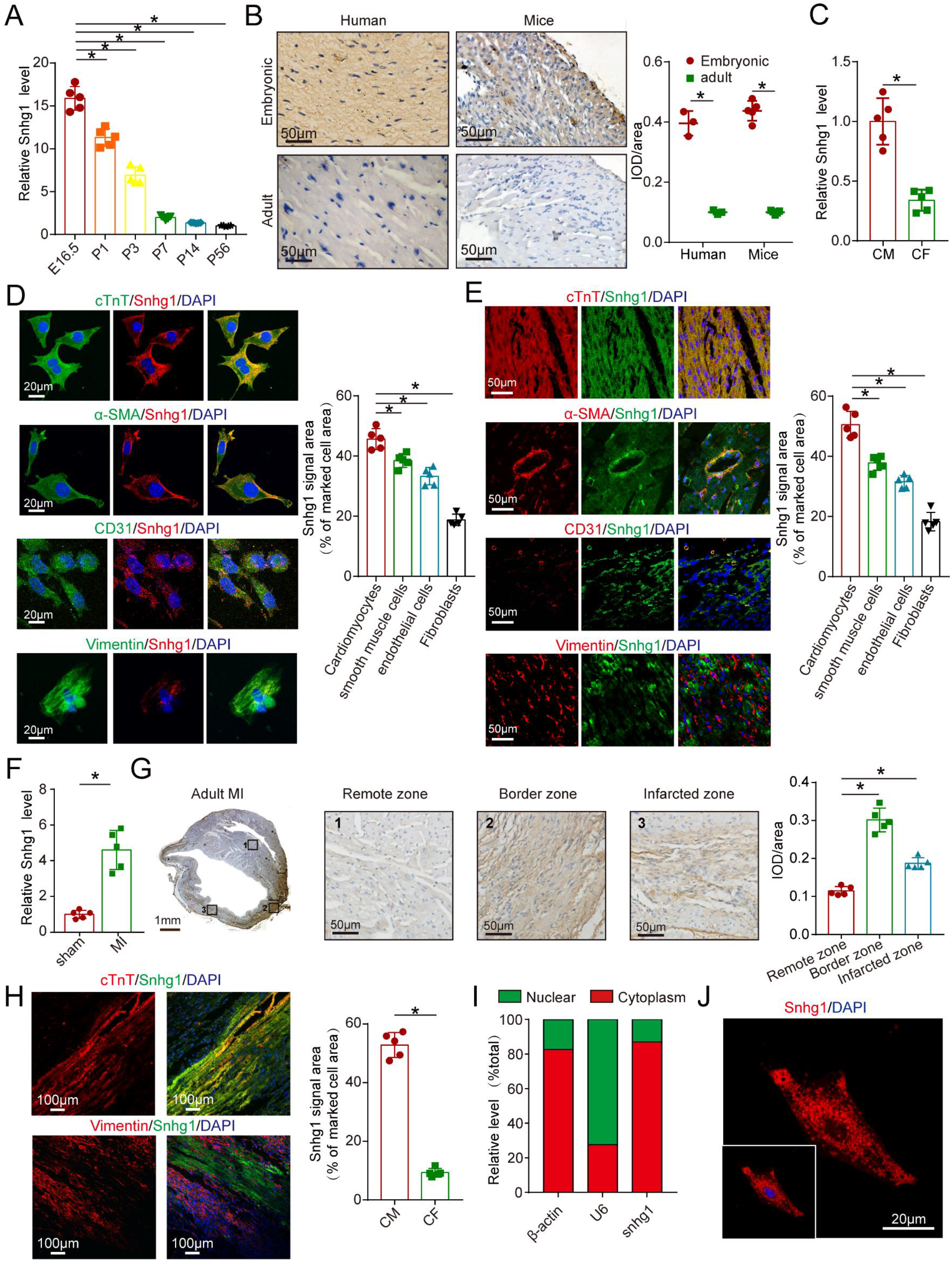
Snhg1 is highly expressed in fetal and MI hearts. (A) Quantification of Snhg1 expression in ventricles harvested from E16.5 to P56 mice by RT-qPCR, n=5. (B) Detection of Snhg1 expression in human and mouse heart tissue by ISH from 3-5 hearts, brown dot cluster indicated Snhg1. (C) RT-qPCR of Snhg1 in isolated neonatal CMs and CFs, n=5. (D-E) Co-staining with neonatal CM, smooth muscle, CF, and endothelial markers to identify cells expressing Snhg1 by using RNA-FISH technique *in vitro* (D) and *in vivo* (E), n=5. (F) Quantification of Snhg1 in sham and 28 days after MI in adult mouse hearts by RT-qPCR, n=5. (G) Detection of Snhg1 expression in adult heart tissue after MI by *ISH*, 1-3 respectively indicated remote zone, border zone and infarcted zone, n=5. (H) Co-staining with CM and CF markers to identify cells expressing Snhg1 in adult hearts after MI by using the *RNA-FISH* technique, n=5. (I) QRT-PCR for the abundance of Snhg1 in either the cytoplasm or nucleus of neonatal CMs. (J) Detection of Snhg1 expression in neonatal CMs by *FISH*.

### Snhg1 promotes P7 CM proliferation

To investigate whether Snhg1 promotes P7 CM proliferation in which most CMs normally exit the cell cycle, we overexpressed Snhg1 by uing Adenovirus vectors(Fig.S2A-C). Overexpression of Snhg1 promoted CM proliferation, as shown by immunostaining for DNA synthesis marker 5-ethynyl-2’-deoxyuridine (EdU) (Fig.2A), mitosis marker phosphorylated Histone H3 (pH3) (Fig.2B), and cytokinesis marker (Aurora B) (Fig.2C). Overexpression of Snhg1 promoted pH3 and Aurora B protein expression in CMs (Fig.S2D). We observed that Snhg1 induced an increase in mononucleated CMs and a decrease in binucleated CMs (Fig.2D). Time-lapse imaging of primary P7 CMs isolated from CM-specific Myh6-mCherry transgenic mice showed that overexpression of Snhg1 induced P7 CMs underwent cell division rather than binucleation in mononucleated CMs (Fig.2E and video.1). Flow cytometry assays revealed that Snhg1 promoted the accumulation of P7 CMs cells in the S and G2/M phases of the cell cycle (Fig.2F).In *vivo*, we overexpressed Snhg1 by delivering adeno-associated virus serotype 9 (AAV9)-mediated constructs(Fig.S3A-C). Overexpression of Snhg1 in P7 mice induced a marked increase in Ki67-, pH3-, and Aurora B-positive CMs (Fig.2G-H and Fig.S3D). EdU incorporation was predominantly observed in mononucleated CMs in the Snhg1-injected mice (Fig.2I), indicating that Snhg1 preferentially promotes mononucleated cells to enter S phase. The CM-specific Myh6-mCherry transgenic mice were further used to confirm the proliferative effect exerted by Snhg1 (Fig.2J-K and Fig.S3E). Collectively, these data suggest that Snhg1 was able to induce P7 CM proliferation and was sufficient for induction of cell cycle re-entry.

**Figure 2.**
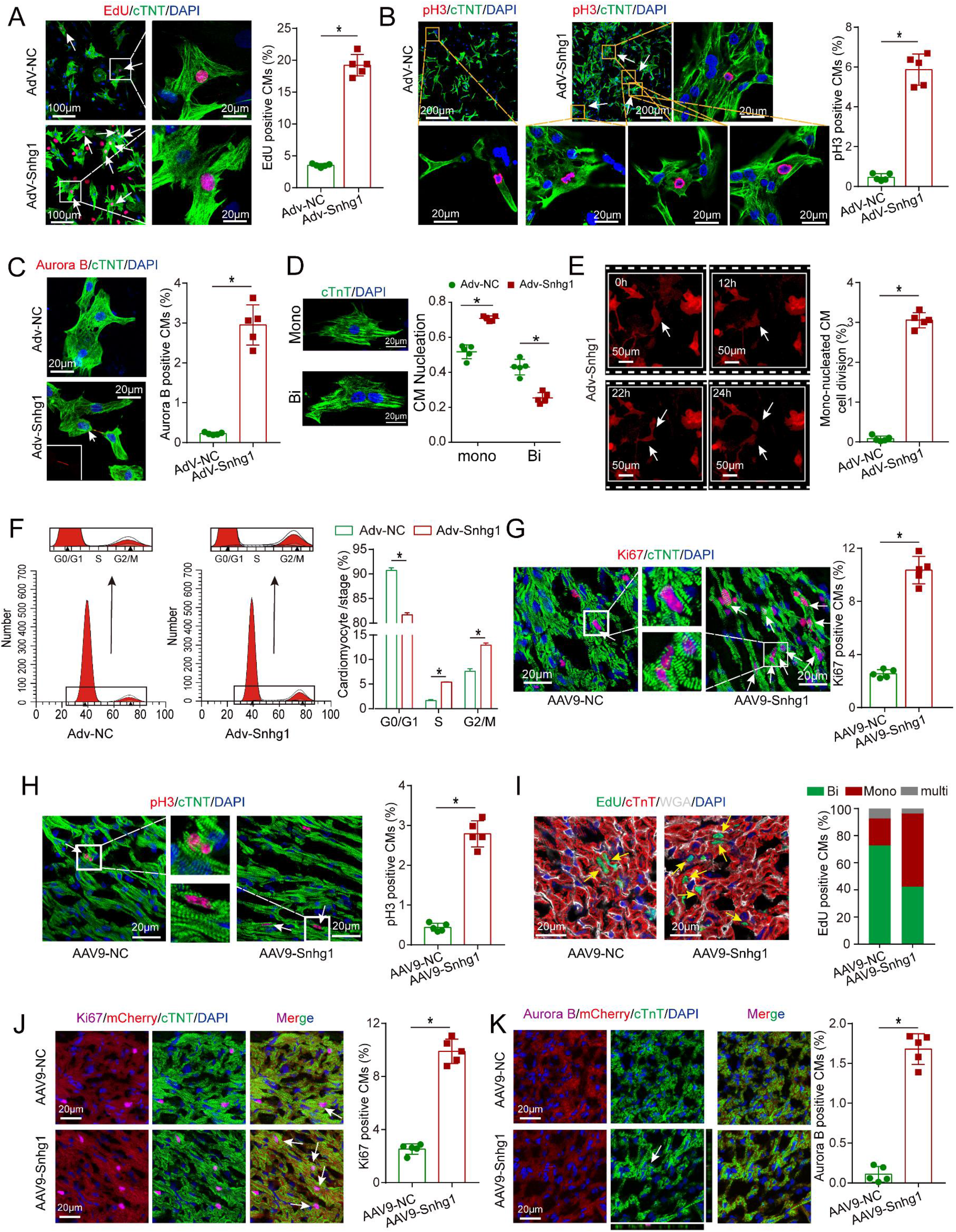
Snhg1 overexpression promotes P7 CM proliferation. (A-C) Immunostaining of EdU, pH3, and Aurora B for P7 CMs transfected with Adv-NC or Adv-Snhg1. CMs were stained with cTnT and nuclei were stained with DAPI. Arrows, positive CMs. n=5. (D) Quantification and representative images of mono-nucleated CMs and binucleation CMs after transfection with Adv-NC or Adv-Snhg1, n=5. (E) Time-lapse imaging of Snhg1-induced cytokinesis in P7 mouse CMs isolated from Myh6-mCherry transgenic mice. Panels are representative of images recorded cell division (see Movie.1), n=5. Arrows, CM underwent cell division. (F) Detectection of cell cycle alterations in P7 CMs transfected with Adv-Snhg1 or Adv-NC by using flow cytometry analysis. (G-H) Immunostaining of Ki67 and pH3 in P7 mouse hearts, n=5. Arrows, positive CMs. (I) EdU incorporation was detected in CM nuclei and cell borders were labeled with wheat germ agglutinin (WGA), n=5. Arrows, positive CMs. (J-K) Immunostaining of Ki67 and Aurora B in P7 Myh6-mCherry transgenic mice, n=5. All CMs were stained with cTnT and nuclei were stained with DAPI. Arrows, positive CMs.

To test the role of Snhg1 in human CMs, we overexpressed Snhg1 in postmitotic 60-day-old human-induced pluripotent stem-cell-derived CMs (hiPS-CMs) and observed an increase in pH3-positive hiPS-CMs (Fig.S4A). Snhg1 had no significance on the proliferation of CFs whether Snhg1 was overexpressed or inhibited, as assessed by EdU incorporation (Fig.S4B-C). Because only 7.2% of cultured CFs were infected with the same adenovirus dose delivered used for CMs (10 MOI) and we had to use 10 times MOI used for CMs on Thy1^+^ CFs to reach a >95% infection efficiency (Fig.S4D).

### Snhg1 induces adult CM proliferation

To investigate the role of Snhg1 on adult CM proliferation, we isolated CMs from adult mouse hearts injected with AAV9-Snhg1 or AAV9-NC (Fig.S5A). As expected, overexpression of Snhg1 in adult CMs resulted in a striking increase in cell cycle activity, a higher ratio of mononucleated CMs, and CM number (Fig.3A-D). We further induced Snhg1 overexpression in adult mouse hearts by intracardiac injection of AAV vectors expressing Snhg1 (Fig.3E-F and Fig.S5B-D). overexpression of Snhg1 increased the ratio of proliferative CMs (Fig.3G-I) and mononucleated CMs (Fig.3J). Stereological analysis further confirmed the increase in CM number in the AAV9-Snhg1 group (Fig.3K and Fig.S6A). All of the above results indicated that overexpression of Snhg1 promoted adult CM proliferation.

**Figure 3.**
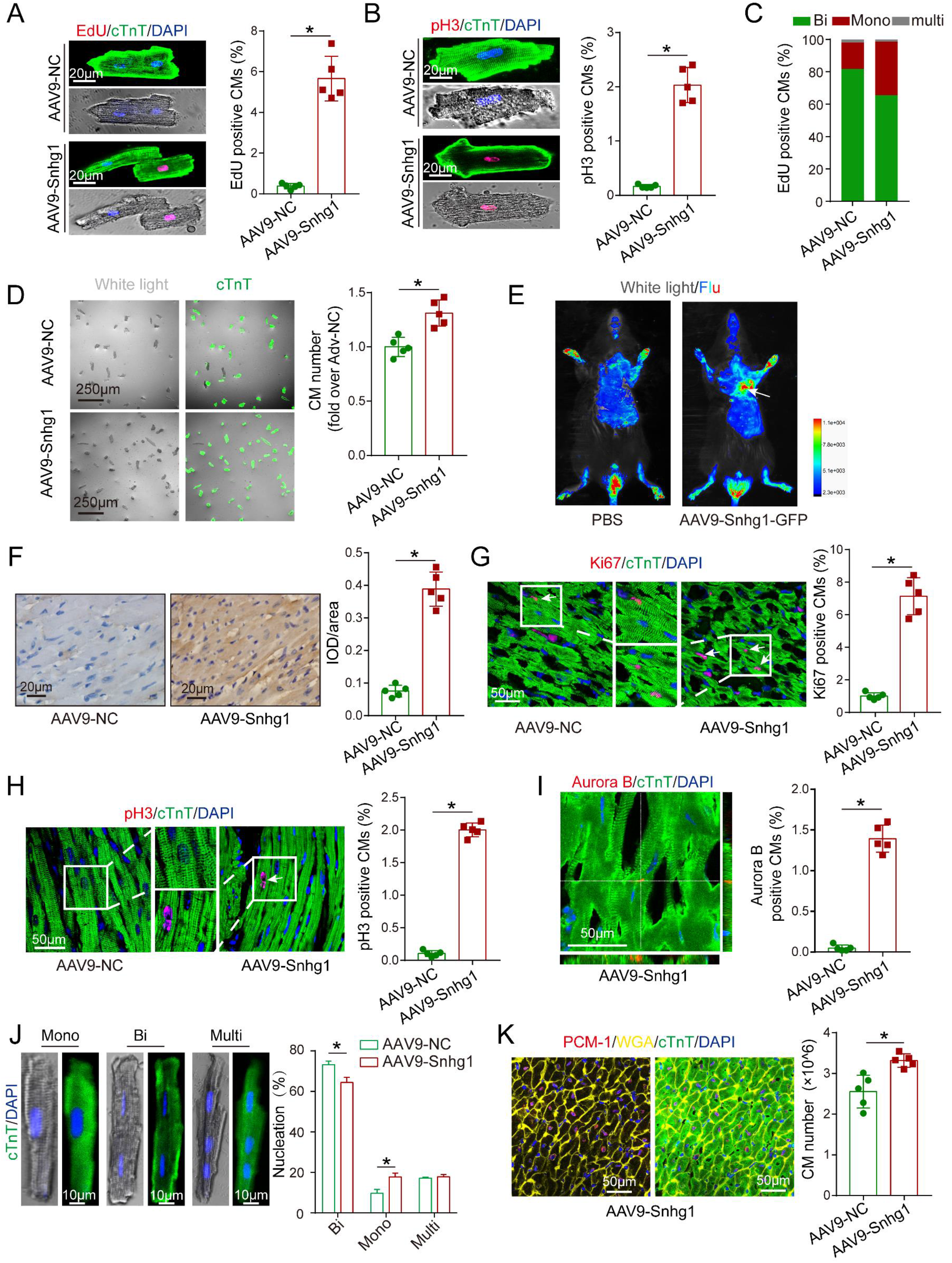
Snhg1 promotes CM proliferation in adults. (A-B) Immunostaining of EdU and pH3 of isolated CMs from adult mouse hearts after transfection with AAV9-Snhg1 and AAV9-NC for 14 days, n=5. (C) EdU incorporation was detected in CM nuclei, n=5. (D) Representative bright field and cTnT immunostaining images of isolated adult CMs from adult mouse hearts after transfection with AAV9-Snhg1 and AAV9-NC for 14 days, and quantification of the CM number, n=5. (E) Representative In vivo bioluminescent images and Bright field captured on day 14 after injection with GFP-labelled AAV9-Snhg1 virus, arrow indicates the heart with GFP fluorescence. (F) *ISH* assays confirmed that Snhg1 was significantly increased in adult hearts injected with AAV9-NC or AAV9-Snhg1. The brown dot cluster indicates Snhg1, n=5. (G-I) Immunostaining of Ki67, pH3 and Aurora B in adult mice hearts, n=5. Arrows, positive CMs. (J) Immunostaining of isolated CM with cTnT and quantification of the nuclei number. For nucleation analysis, approximately 1 × 10^3^ CMs were counted per sample, n=5. (K) Stereological analysis revealed the number of CMs in adult mice, n=5.

### Snhg1 induces marked cardiac regeneration and improved functional recovery after infarction

To determine whether Snhg1 improved in cardiac repair, we delivered the AAV9-Snhg1 to peri-infarcted area in adult mice after MI induced by permanent ligation of the LAD coronary artery (Fig.S7A-B). *ISH* analysis confirmed that intracardiac injection of AAV9-Snhg1 significantly increased Snhg1 expression in both sham operated and MI hearts (Fig.4A). Overexpression of Snhg1 increased the number of Ki67-, EdU- and pH3-positive cells in the peri-infarct areas (Fig.S7C and Fig.4B-C). These results were further confirmed in CM-specific Myh6-mCherry transgenic mice (Fig.S7D). Overexpression of Snhg1 resulted in an decrease in myocyte apoptosis in the infarct border zone Fig.4D) and an increase in blood vessel density in the peri-infarct zone (Fig.4E-F). Masson’s trichrome staining clearly showed that the infarct size was significantly reduced in Snhg1-overexpressing hearts (Fig.4G). As evaluated by echocardiography, Snhg1 overexpression improved the cardiac function in mouse hearts at 21 and 28 days after MI (Fig.4H). The above results were confirmed by using the ischemia-reperfusion (I/R) injury mouse model (Fig.S8A-G). Taken together, Snhg1 improved cardiac function and reduced fibrotic area, which were probably attributed to the increase in CM proliferation and angiogenesis and reduction in CM apoptosis.

**Figure 4.**
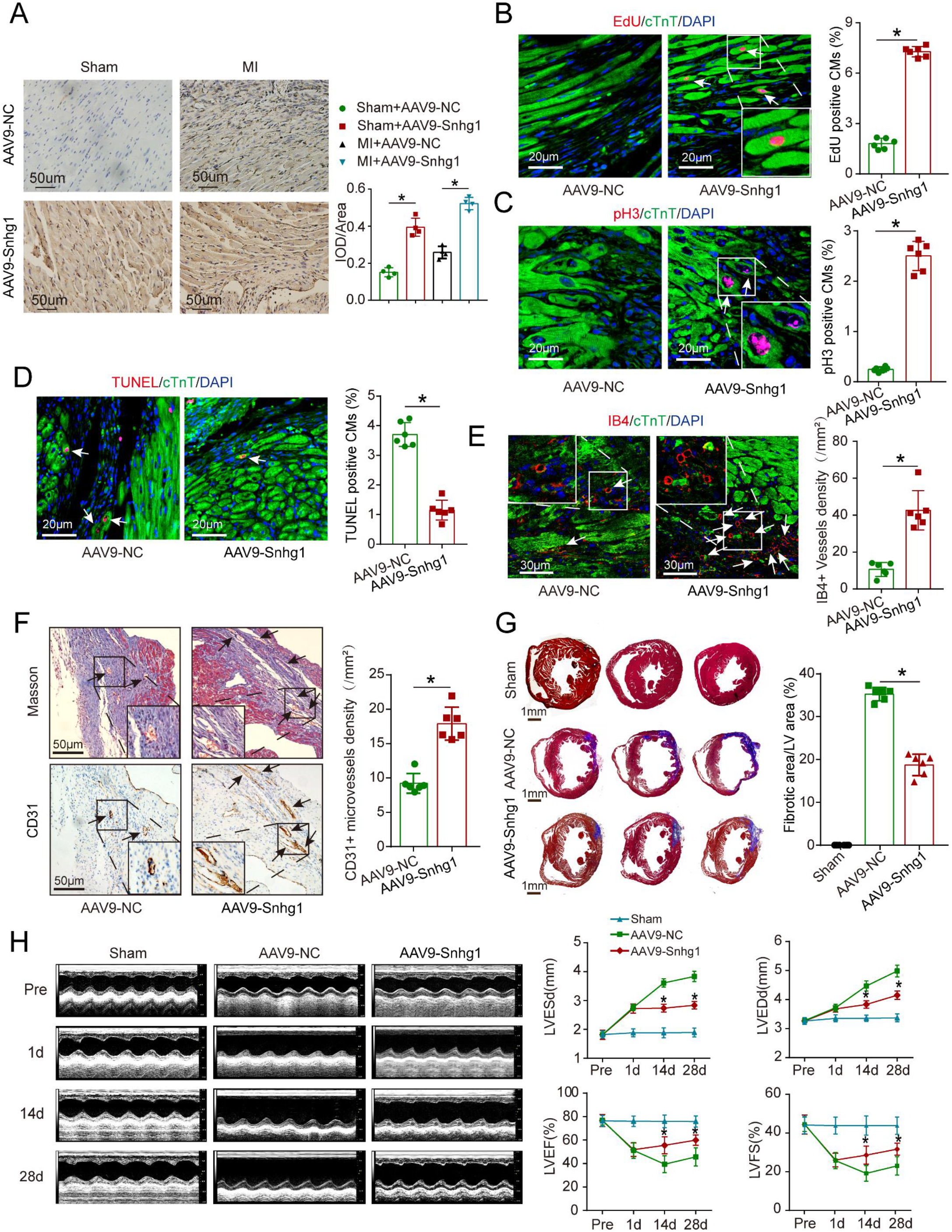
Snhg1 improves adult cardiac function post-MI. (A) Detection of Snhg1 expression in sham operated and infarcted adult mouse hearts by using ISH analysis, n=4. (B-C) Immunofluorescence of EdU and pH3, TUNEL and IB4 in adult mouse hearts at 14 days after MI, n=6. Arrows, positive CMs. (D-E) Immunofluorescence of TUNEL and IB4 in adult mouse hearts at 7 days after MI, n=6. Arrows, positive cells or vessels. (F) Immunohistochemistry for CD31 in adult mice at 14 days after MI, n=6. (G) Masson’s trichrome-stained heart sections in adult mice at 28 days after MI and quantification of infarct size, n=6. (H) Echocardiography analysis of adult mouse hearts at 1,14 and 28days after MI, n=10.

### Snhg1 depletion impaired cardiac regeneration in neonatal mouse

We further knocked down Snhg1 in P1 mouse CMs to determine whether inhibition of Snhg1 impairs neonatal cardiac regeneration(Fig.S9A). Depletion of Snhg1 decreased the ratio of proliferative CMs and induced CMs were accumulated at the G0/G1 phase of cell cycle(Fig.5A-D), and also decreased pH3 and Aurora B protein expression(Fig.S9B). At 5-7 days after intracardiac injection of Adv-si-Snhg1, Snhg1 was decreased in neonatal mouse hearts (Fig.S10A-B). Inhibition of Snhg1 resulted in a decrease in CM proliferation (Fig.5E, Fig.S10C-D). Snhg1 depletion by using Adv-si-Snhg1 significantly decreased the Snhg1 level in both sham operated heart and MI neonatal hearts (Fig.5F). At 5-7 days post-MI, systolic function was significantly impaired (Fig.S10F and Fig.5G) and the fibrosis area was greater in the Adv-si-Snhg1 group (Fig.5H). To investigate whether endogenous Snhg1 is a critical regulator of CM proliferation, we used CRISPR-Cas9 knock-in mice to generate loss-of-function mutations in Snhg1 genes (Fig.S11A-F). We observed a significant decrease in CM proliferation in these Snhg1-deficient mice (Fig.S11G-H). Collectively, these results suggest that downregulation of Snhg1 impaired neonatal cardiac regeneration.

**Figure 5.**
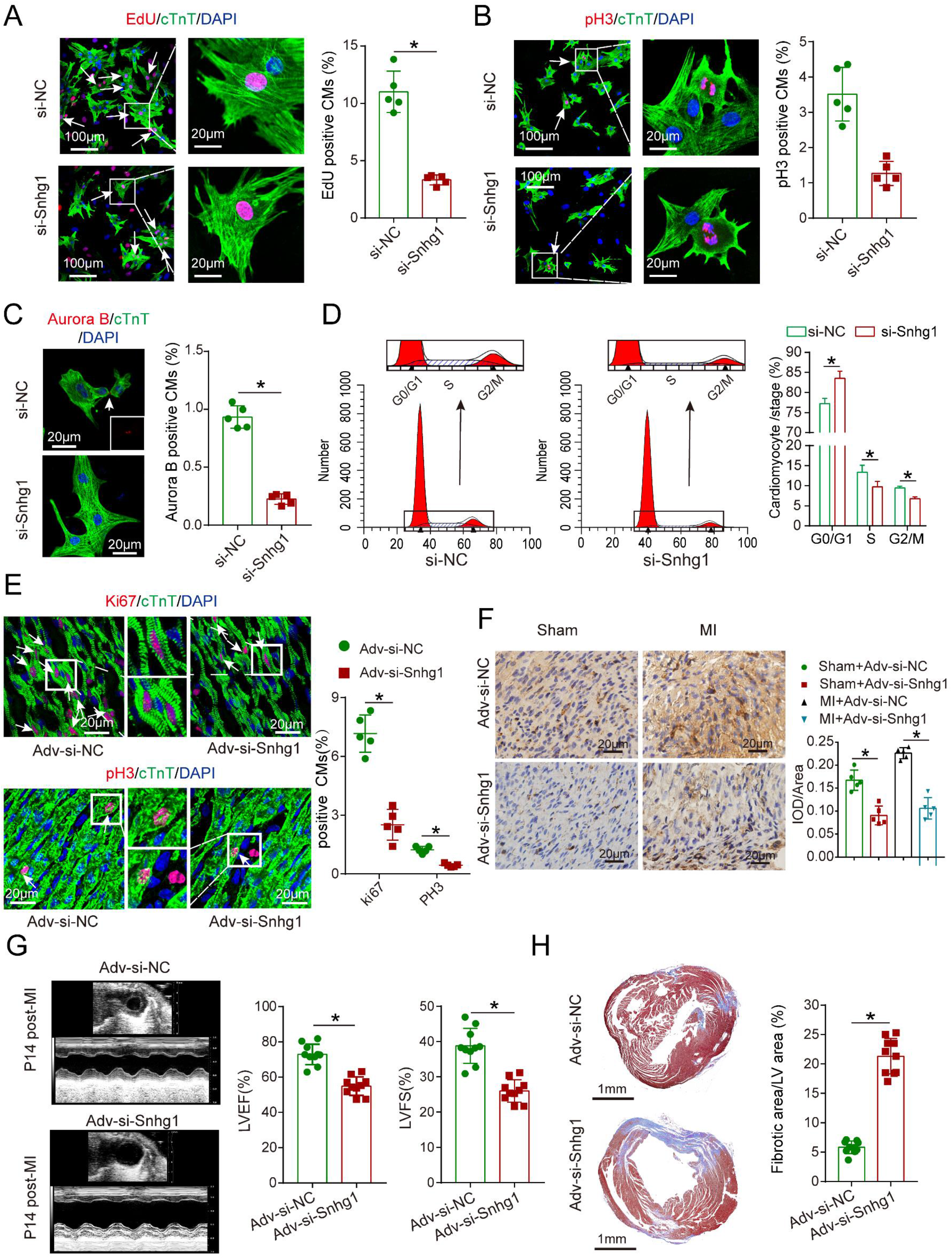
Loss of Snhg1 inhibits neonatal CM proliferation. (A-C) Representative images and quantification of P1 CMs positive for EdU, pH3, and Aurora B, n=5. Arrows, positive CMs. (D) Detectection of cell cycle alterations in P7 CMs using Flow cytometry analysis. (E) Immunofluorescence of Ki67 and pH3 in neonatal hearts, n=5. Arrows, positive CMs. (F) Detection of Snhg1 expression in sham operated and infarcted neonatal mouse hearts by using ISH analysis, n=5. (G-H) Echocardiography and Masson’s trichrome-stained heart sections of neonatal mouse hearts detected by echocardiography in neonatal mice at 5-7 days after MI and quantification of LVEF and LVFS, n=10.

### Snhg1 regulates CM proliferation through the PTEN/PI3K-Akt/c-Myc pathway

To explore how Snhg1 mediated cardiac regeneration, we performed next-generation RNA sequencing (RNA-seq) in P7 mice by overexpressing Snhg1. The transcriptome analysis identified 4029 genes were significantly upregulated and 1449 genes were significantly downregulated(Fig.6A and Fig.S12A). The upregulated genes were associated with cell development or differentiation(Fig.S12B). The top enriched pathways of differentially upregulated genes included cell cycle, PI3K-Akt and Hippo signaling pathways (Fig.6B), which played vital roles in CM regeneration. The differentially expressed genes enriched in the cell cycle, PI3K-Akt and Hippo signaling pathways overlapped (Fig.6C-D and Fig.S12C). These findings suggest that Snhg1 plays a prominent role in mediating CM proliferation.

**Figure 6.**
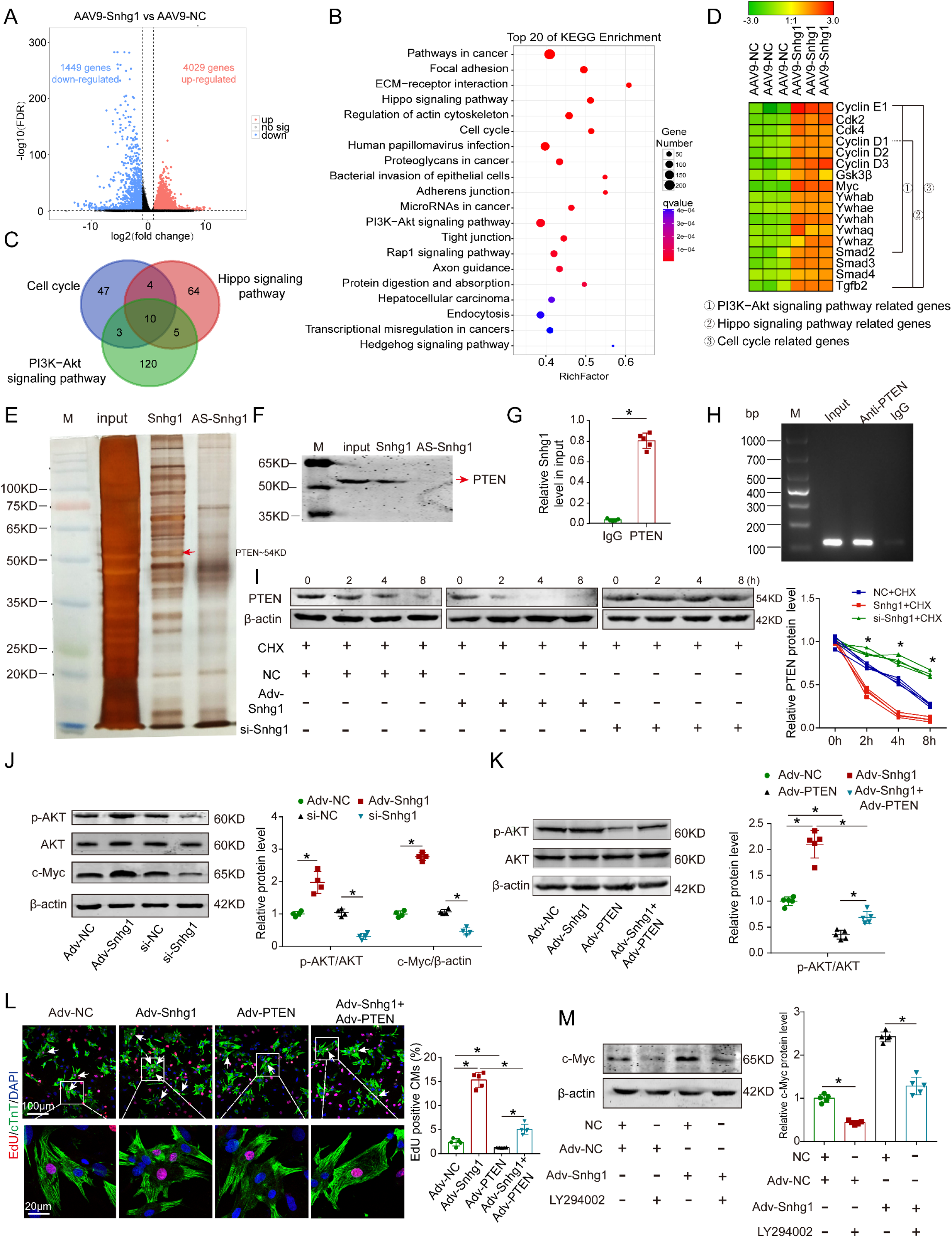
Snhg1 regulates CM proliferation via the PTEN/Akt/c-Myc pathway. (A)Volcano plot displaying differential expressed genes in P7 hearts. The red dots represent the up-regulated expressed genes and blue dots represent the downregulated expressed genes. (B) The top 20 KEGG pathway enrichment scatter plot of up-regulated genes. x-axis indicates the rich factor, y-axis specifies the KEGG pathways. (C)Venn diagram showing the number of overlapping genes enriched in KEGG pathway genes from the cell cycle, PI3K-Akt, and Hippo pathways. (D) Representative gene modules of overlapping genes. (E) Silver-stained SDS-PAGE gel of proteins immunoprecipitated by RNA pull-down assay of Snhg1 or its antisense RNA (AS-Snhg1). Arrow indicated the indentified PTEN protein. (F) PTEN protein was assayed by Western blotting. (G-H) RIP were performed using an antibody against PTEN or negative IgG. Purified RNA was used for RT-qPCR analysis, and enrichment of Snhg1 was normalized to the input, n=5. (I) PTEN protein levels were measured by Western blotting in isolated CMs. CHX was used to block protein synthesis, n=4. (J)Western blotting analysis of p-Akt, Akt, and c-Myc protein in isolated CMs with Snhg1 overexpression or depletion, n=4. (K) Western blotting analysis of p-Akt, and Akt protein levels. (L) immunofluorescence of EdU in isolated CMs, n=5. Arrows, positive CMs. (M) Western blotting analysis of c-Myc protein levels in isolated CMs.

We further performed RNA pull-down assays in isolated P7 CMs to identify the protein that interacts with Snhg1(Fig.6E) and Mass spectrometry (MS) identified the phosphatase and tensin homolog (PTEN) as the possible protein that might interacts with Snhg1(Fig.S13A). Western blotting and RNA immunoprecipitation (RIP) verify the interaction between Snhg1 and PTEN (Fig.6F-H). Overexpression of Snhg1 decreased the PTEN protein levels, and knockdown of Snhg1 showed the opposite effect on the PTEN protein (Fig.S13B). by using the protein synthesis inhibitor cycloheximide (CHX), we further found that Snhg1 decreased PTEN protein stability(Fig.6I). We further determined whether Snhg1 regulates the PI3K-Akt/c-Myc pathway via PTEN. Overexpression of Snhg1 increased the phosphorylated Akt (p-Akt) and c-Myc protein levels, while depletion of Snhg1 showed the opposite effects (Fig.6J). Rescue of PTEN counteracted the increase in p-Akt protein levels and proliferative effect by Snhg1 overexpression (Fig.6K-L and Fig.S3C-E). Inhibition of Akt phosphorylation by using LY294002 abrogated the elevated expression of c-Myc proteins induced by Snhg1 (Fig.6M and Fig.S13F). Then we explored how Akt regulates c-Myc. Depletion of Snhg1 or inhibition of Akt phosphorylation decreased both the phosphorylation of GSK3β and the c-Myc protein levels(Fig.S13F), and the decreased of c-Myc protein by LY294002 could be partly restored by inhibition of GSK3β(Fig.S13G). Thus, Snhg1 induces Akt phosphorylation by decreasing the stability of PTEN, leading to the inhibition of GSK3β, thereby preventing the proteasome-mediated proteolysis of c-Myc. All of the above results demonstrated that Snhg1 regulates CM proliferation via the PTEN/Akt/c-Myc pathway.

### c-Myc/Snhg1 forms a positive feedback loop

We next investigated the upstream regulator of Snhg1. By using the Jaspar database, we predicted that c-Myc has two potential binding sites in Snhg1 promoter (Fig.7A). DNA pull-down assays following by Western blotting revealed that c-Myc can bind to the promoter region of Snhg1 (Fig.7B). ChIP-qPCR analysis revealed that c-Myc directly binds to site 1 of the Snhg1 promoter (Fig.7C). We next investigated whether c-Myc regulates Snhg1 in CMs. Overexpression of c-Myc increased the Snhg1 levels, whereas suppression of c-Myc had the opposite effect (Fig.7D). Moreover, inhibition of c-Myc counteracted the proliferative effect by Snhg1 overexpression(Fig.7E-F). Overexpression of c-Myc increased p-Akt expression and decreased PTEN expression in CMs while suppression of c-Myc showed the opposite effects (Fig.7G). These data indicated that c-Myc/Snhg1 form a positive feedback loop in CMs. We further explored whether this positive feedback loop is self-limiting as time progresses. We found that overexpression of Snhg1 increased the Snhg1 level started from 2 days and remained stable for at least 7 days, while the levels of its downstream proteins c-Myc declined at the 6^th^ day and became stable thereafter (Fig.7H-I). Luciferase assay revealed that as the level of c-Myc protein increases, the inhibition of c-Myc promoter activity increases and gradually reaches a steady state (Fig.7J), indicating that c-Myc drives an autoregulatory feedback loop.

**Figure 7.**
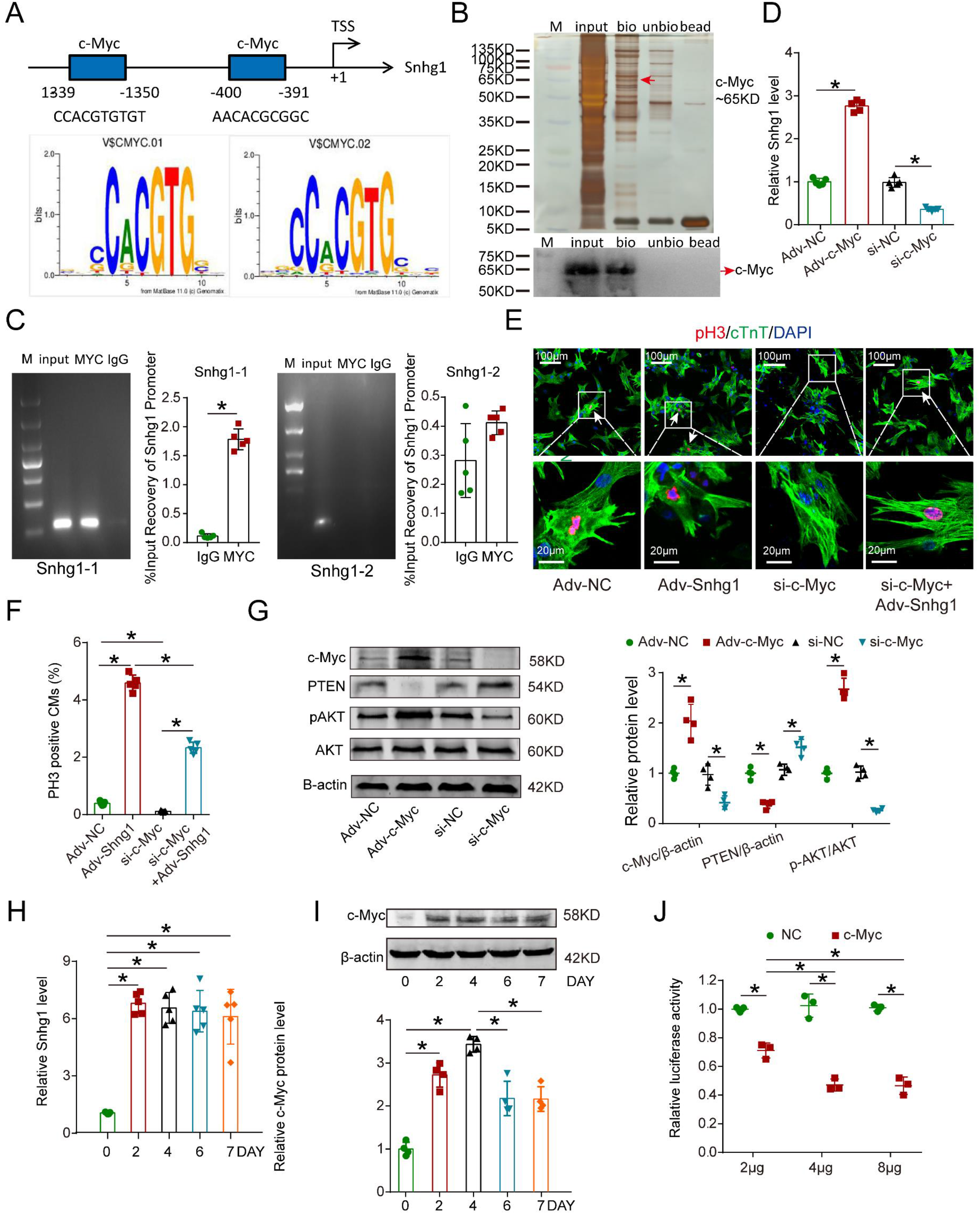
c-Myc upregulates Snhg1 expression by binding to its promoter region. (A) The two predicted c-Myc binding regions and sequences in the promoter region of Snhg1. (B) Silver-stained SDS-PAGE gel of proteins immunoprecipitated by DNA pull-down assay of biotinylated or unbiotinylated probe targeting Snhg1. Arrow indicates region of the gel excised for MS determination. c-Myc protein was assayed by Western blotting. (C) Image of agarose electrophoresis showing ChIP-qPCR using anti-c-Myc or anti-IgG antibodies was performed to access binding between Snhg1 and c-Myc. Purified RNA was used for RT-qPCR, and enrichment of Snhg1 was normalized to the input, n=5. (D) RT-qPCR of Snhg1 levels in isolated P7 CMs of different groups, n=5. (E-F) Immunofluorescence of pH3 of P7 CMs transfected with Adv-NC, Adv-Snhg1, si-c-Myc, or si-c-Myc + Adv-Snhg1 in P7 CMs, n=5. Arrows, positive CMs. (E) Western blotting analysis of c-Myc, PTEN, p-Akt, and Akt protein levels in isolated CMs overexpressed or depleted with c-Myc, n=4. (H) RT-qPCR of Snhg1 levels in isolated P7 CMs transfected with Adv-Snhg1 for 0, 2, 4, 6, and 7 days, n=5. (I) Western blotting analysis of c-Myc protein levels in isolated P7 CMs transfected with Adv-Snhg1 for 0, 2, 4, 6, and 7 days, n=4. (J) P7 CMs transfected with the c-Myc promoter (PGL3-c-Myc plasmid) and different concentrations of the c-Myc protein. The background luciferase activity (empty vector) was subtracted from all data, n=3.

To detect whether c-Myc-bound Snhg1 promoter is key for inducing CM proliferation, Snhg1 promoter regions with or without the c-Myc-binding sequence mutation vectors (MU) were constructed (Fig.8A). After the c-Myc-binding sequence was mutated, overexpression of Snhg1 no longer increased snhg1levels(Fig.8B). In addition, Overexpression of Snhg1 did not affected the PTEN, p-Akt and c-Myc expression(Fig.8C) and CM proliferation (Fig.8D-E) when the c-Myc-binding sequence was mutated. To directly visualize CM divisions, we conducted time-lapse imaging of P7 CMs labeled with the fluorescent mitochondrial dye tetramethylrhodamine ethyl ester. We observed that Snhg1 overexpression-derived CMs underwent cytokinesis (Fig.8F and video.2), while the CMs underwent karyokinesis as was observed after the c-Myc-binding sequence was mutated(Fig.8G and Video.3), and no cell division was observed in the NC group (Fig.8H and Video.4). These results revealed that the Snhg1 promoter without the c-Myc-binding region could not robust induce CM proliferation.

**Figure 8.**
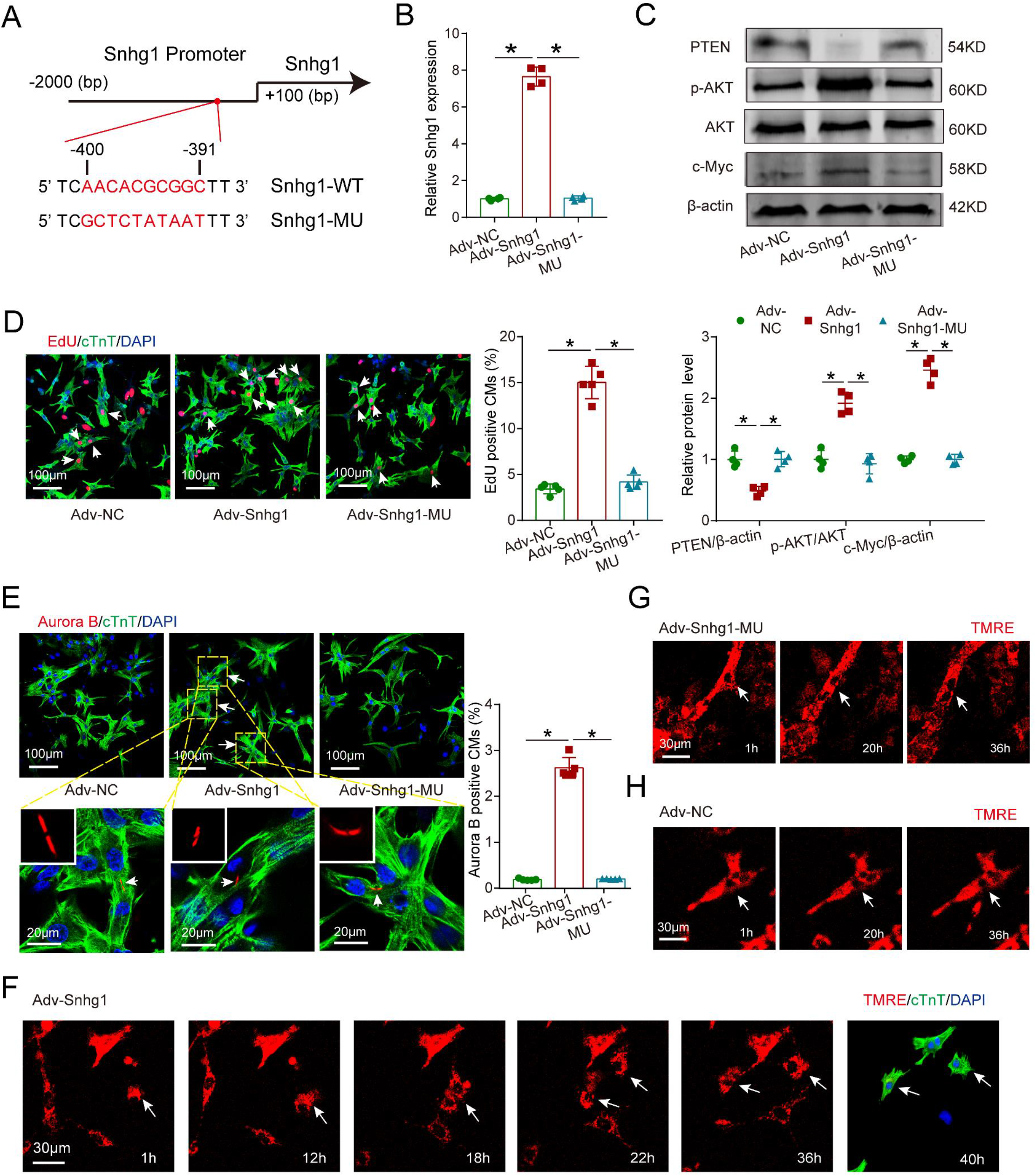
Mutant of Snhg1 inhibits Snhg1-induced CM proliferation. (A) Construction of Snhg1-WT and Snhg1-MU promoter sequences. (B-C) RT-qRCR of Snhg1 levels and western blotting analysis of PTEN, PI3K, p-Akt, Akt, and c-Myc protein levels in isolated CMs, n=4. (D-E) EdU and Aurora B immunofluorescence staining in P7 CMs, n=5. Arrows, positive CMs. (F-H) Representative images from time-lapse videos of P7 CMs transfected with Adv-Snhg1 (F), Adv-Snhg1-MU (G) or Adv-NC (H)

## Discussion

Our study demonstrated that the lncRNA Snhg1 drives a positive feedback loop that efficiently induces CM proliferation and subsequent improvement in functional recovery after MI. Mechanistically, Snhg1 formed a positive feedback loop with c-Myc to drive a self-reinforcing circuit that sustained activation of the PTEN/PI3K-Akt/c-Myc pathway, resulting in continuous cell cycle re-entry, which may contribute to CM regeneration (Fig.S14). Our findings reveal that Snhg1 can stimulate cardiac regeneration and may represent a promising therapeutic target for heart failure.

Increasing evidence has shown that the adult mammalian heart can undergo some self-renewal at a low level (Bergmann *et al*, 2009). However, few or no CMs undergo cytokinesis to form new daughter cells (Senyo *et al*, 2013). Our study provides important insights into the postnatal role of Snhg1 in controlling CM cytokinesis and cardiac regeneration. In the present study, we evaluated proliferative effects not only performing immunostaining of cell cycle marker (Ki67 and EdU) analysis, classical cell mitosis and cytokinesis marker (pH3 and Aurora B kinase) analysis, but also used flow cytometry assays to revealed the cell cycle progression. In addition, we used CM-specific Myh6-mCherry transgenic mice to confirme the effecf of Snhg1 on CM proliferation. Moreover, we performed an *in vitro* cytokinesis assessment to observe incidence of cytokinesis in real-time by using time-lapse fluorescence microscopy. Strikingly, echocardiography and Masson’s trichrome staining supported the involvement of Snhg1 in enhancing heart regeneration and functional improvement in both neonatal and adult mouse hearts post-MI. Thus, our results confirm that Snhg1 can stimulate CM proliferation and improve cardiac function post-MI.

By using RNA sequencing, we found that Snhg1 mainly induced gene expression related to cell cycle, PI3K-Akt and Hippo signaling pathways, which are involved in cardiac regeneration. PI3K-Akt signaling played important roles in controlling cell division.(Amit & Ana C, 2007) Activating PI3K-Akt signaling, either directly or by interfering with its upstream regulators, likely represents an effective and widely applicable heart regeneration strategy. Our result suggest that Snhg1 is a key upstream regulator of PTEN that modulates Akt phosphorylation by activating the PI3K-Akt pathway to control the commitment to CM proliferation. In addition, Snhg1 promoted the proliferation of endothelial cells. Activation of the PI3K/AKT pathway could increase VEGF secretion and modulate the expression of other angiogenic factors to promote endothelial cell proliferation (Facchini *et al*, 1997; Hamada *et al*, 2005). Thus, we speculated that Snhg1 promoted endothelial cell proliferation by activating the PI3K-AKT pathway.

We further found that Snhg1 forms a positive feedback loop with c-Myc to sustain activation of PI3K-Akt signaling, which increases CM cytokinesis. c-Myc is a master transcription factor that regulates genes essential for cell proliferation, differentiation, and apoptosis (Dang, 1999). Our result revealed that c-Myc functioned as a transcription factor by binding to the promoter regions of Snhg1 and upregulating the Snhg1 expression level, at the same time, c-Myc was a downstream target of PTEN/PI3K-Akt signaling, revealing a positive feedback loop between c-Myc and Snhg1. Positive feedback is a key mechanism ensuring that a cell is fully committed to cell division (Santos *et al.*, 2012). If the positive feedback loop is perturbed, mitosis become longer and more variable, causing cells to die during mitosis or soon after without reaching a second round of cell division (Araujo *et al*, 2016). In this study, we identified the critical interplay between Snhg1 and PI3K/Akt signaling, which drives the onset of a positive feedback loop through the transcriptional activation of the c-Myc genes. Our findings potentiated the ability of Snhg1 to promote CM cytokinesis.

Although our data showed that Snhg1 is promising for cardiac regeneration, there are some potential limitations. Positive feedback loops tend to lead to instability, and a lack of control may cause ongoing cellular division. Because excessive or unwanted cell division may increase the risk of tumorigenesis, the delivery of Snhg1 must be carefully controlled. Transient *in vivo* delivery and organ-specific delivery systems may reduce the risks associated with tumorigenesis. Although we did not observe any cardiac tumors in mice treated with Snhg1, the potential for ectopic proliferation requires careful and specific delivery in future clinical development. Another risk is activation of the immune response associated with viral delivery systems. The immune system may block the virus from delivering the gene to the heart, impacting the effectiveness of Snhg1. Given the pleiotropic effects of lncRNAs, effective translation of Snhg1-based treatments into a successful clinical therapy may require a deeper understanding of the underlying molecular and cellular mechanisms driving Snhg1-induced CM proliferation. Moreover, these results must be further validated in large animal models for cardiac regeneration post-MI or other ischemic injury.

In conclusion, our data showed that Snhg1 forms a positive feedback loop with c-Myc to sustain the activation of PI3K/Akt signaling, which effectively elicits CM cytokinesis and improves function post-MI. These findings suggest that Snhg1 might represent as a powerful regenerative approach for treating heart failure.

## Acknowledgments

We thank Dr. Jinzhu Duan (Guangzhou Women and Children’s Medical Center, Chin) for kindly providing the Myh6-mCherry transgenic mice.

## Data availability

The raw/processed data required to reproduce these findings are available from the corresponding author on reasonable request.

## Legends for videos

**Video 1.** Time-lapse movie of cell divisions of P7 mouse CMs isolated from Myh6-mCherry transgenic mice overexpressing Snhg1.

**Video 2–4.** Time-lapse videos of P7 CMs transfected with Adv-Snhg1 (Video. 2), Adv-Snhg1-MU (Video.3) or Adv-NC (Video. 4).

## Detailed Methods

### Human tissue samples

Human left ventricular tissue samples were obtained as described previously.^1^ Briefly, adult myocardial tissue samples were obtained by endomyocardial biopsy from patients suffering from myocardial deposition disease based on arrhythmia and echocardiographic changes. The pathological findings showed no evidence of myocardial disease or functional abnormalities. Embryonic human myocardial samples were obtained after elective termination of pregnancy for nonmedical reasons. The present study conformed to the principles of the Declaration of Helsinki. The study protocol was approved by the Nanfang Hospital ethics committee, and written informed consent was obtained from all subjects.

### Human iPSC-CM cultures

CMs derived from iPSCs were purchased from Cellapy (Beijing, China) and were cultured according to the manufacturer’s instructions. The cells were allowed to adhere for 48 h before maintenance medium exchange and fresh medium was replaced every other day and then for transfection experiments.

### Isolation and culture of neonatal CFs

CFs were isolated from mice as previously described^3^. Excised hearts were rinsed in cold Hank’s balanced salt solution (HBSS), minced, and digested with type II collagenase and pancreatin at 37°C for 15 min. The first digestion was discarded. A second digestion was performed, and the collagenase medium containing CFs was collected, centrifuged, and resuspended in DMEM with 10% FBS, 100 U/mL penicillin, and 100 mg/mL streptomycin. Digestion was repeated 5-6 times until the digestion fluid became clear. Cells were plated in 60-mm dishes and allowed to attach for 60 min, and the media was then changed to remove CMs and endothelial cells. Isolated CFs were washed twice with PBS and cultured for transfection experiments.^1^

### EdU administration *in vitro* and *in vivo*

In the *in vitro* experiment, for the 1-day-old and 7-day-old CMs, after transfection for 24 h, the culture medium was replaced by fresh medium for 24 h. That is, 48 h after the cultured cells were seeded, 10μM EdU was added for 24 h. Cells were fixed at 72 h after seeding and processed for immunofluorescence. Adult CMs were isolated from adult mice that were transferred AAV9-mediated Snhg1 or NC. EdU was administered intraperitoneally (500 mg per animal) at 1, 3, 5, 7, 9, 11, 13 days after transfection. CMs were isolated and processed for immunofluorescence 14 days after transfection.

For the *in vivo* experiment, the P1 mice received EdU (50 μg/g) by intraperitoneal daily injections at P5 and P6. The hearts of the injected mice were collected 7 days after transfection. In P7 and adult mice, EdU was administered intraperitoneally (50 μg/g) at 12, 13 days after transfection, and the hearts were collected 14 days after transfection. In adult mice, EdU was administered intraperitoneally (500 mg per animal) at 1, 3, 5, 7, 9, 11, 13 days after transfection, and the hearts were collected 14 days after transfection.

### Immunofluorescence analysis

For *in vitro* cultured CMs, the culture medium was washed with PBS. The cells were fixed with 4% paraformaldehyde (Leagene), permeabilized with 0.2% Triton X-100 PBS, and blocked with PBS containing 1% BSA. The cells or slides were incubated with primary antibodies, including cardiac troponin T (Abcam, ab33589), PCM1 (Abcam, ab72443), vimentin (Abcam, ab8978), Ki67(Abcam, ab15580), pH3 (Abcam, ab170904), Aurora B (Abcam, ab2254), and Anillin (Santa Cruz Biotechnology, sc-271814) for 2 h at room temperature, followed by incubation with goat anti-mouse IgG/Alexa Fluor 488 or goat anti-rabbit IgG/Alexa Fluor 555 secondary antibodies (Biosynthesis, bs-0296GA488, bs-0295G-AF555) for 1 h at room temperature. The cells or slides were washed and incubated with DAPI (BioWorld, St. Louis Park, MN, USA). CM borders were defined by staining of tissue with WGA conjugated to Alexa Fluor 555 (Invitrogen) in PBS. Image acquisition was performed with an LSM 880 confocal microscope (Zeiss, Oberkochen, Germany). The Click-iT® EdU Imaging Kits (Life Technologies, USA) to detect EdU incorporation were used according to the manufacturer’s instructions. Finally, cells were stained with DAPI.^5–7^

For the *in vivo* experiments, formalin-fixed tissue slides were deparaffinized, and antigen retrieval was performed by microwaving the slides in citrate buffer (0.1 mol/L, pH 6.0) for 14 min. When indicated, the cells or slides immunostained with EdU were further processed using a Click-iT EdU Alexa Fluor 555 Imaging Kit (Invitrogen) to reveal EdU incorporation according to the manufacturer’s instructions. Slides were processed for immunofluorescence as described above for the cultured cells.

### *In situ* hybridization (ISH)

ISH was performed utilizing the Panomics QuantiGene ViewRNA ISH tissue assay (Affymetrix, Santa Clara, CA, USA) as previously described.^1^ Mouse hearts were fixed in 10% formaldehyde and embedded in paraffin. Five-micron sections were cut, deparaffinized, boiled in pretreatment solution, and digested with proteinase K. Heart sections were hybridized for 3h at 37°C with a custom designed probe against Snhg1 (Affymetrix). Bound probes were amplified according to the protocol from Panomics using PreAmp and Amp molecules. Working label probe oligonucleotides conjugated to 6-AP were added. AP-Enhancer solution was added to each tissue section after washing. Working label probe oligonucleotides conjugated to 1-AP were added. Slides were counterstained with hematoxylin. Images were acquired with a Nikon Eclipse TE2000-S microscope (Nikon, Tokyo, Japan).

### RNA Fluorescent in Situ Hybridization (RNA-FISH)

Isolated CMs grown on coverslips were fixed in 4% paraformaldehyde, and frozen sections (5 μm) of hearts were fixed with 95% ethanol. Then, the cells or the tissue slides were washed 3 times with PBS. The samples were permeabilized in 0.2% Triton X-100 PBS and washed with PBS for 3 times. Then the samples were refixed with 4% paraformaldehyde for 10 min and dehydrated through sequential 5-minute incubations in ethanol (70%, 80%, 95%, 100%). The samples were incubated with prehybridization solution for 30 min and hybridized with hybridization solution and a labeled Snhg1 probe overnight at 42°C. The sequences of Snhg1 probe were: 5’-AAAACGTGTTATTTGTAAAATTGAACAGGCCTGGCTCCAAAGTGTAAA-3’. Next, the samples were washed with 50% formamide/2×SSC, 0.1% NP40/1×SSC, 0.5×SSC and 0.2×SSC and blocked with PBS containing 1% BSA. The samples were incubated with primary antibodies, including cardiac troponin T (Abcam), α-SMA (Abcam), CD31 (Abcam), and vimentin (Abcam) for 2 h at room temperature, followed by incubation with goat anti-rabbit IgG/Alexa Fluor 594 secondary antibodies (Biosynthesis) for 1 h at room temperature. The cells or slides were washed and incubated with DAPI. Image acquisition was performed with an LSM 880 confocal microscope.

### Stereological analysis

Left ventricles (including the septum) were sampled as previously described.^8,9^ Briefly, heart tissues were embedded in 8% gelatin, and isectors were used to obtain an isotropic, uniform, random alignment of the samples with a maximum diameter of 4 mm. These isectors were used for stereological analysis. An anti-PCM1 antibody was applied to label CM nuclei. WGA was added to identify the cell borders. A minimum of 1-4 isectors were stained, and a minimum of 200 nuclei per animal were counted (nearly 2% of the area of the region of interest). CMs were cut along their longitudinal axis to determine the number of nuclei per cell. The two-step NVⅩ vancomycin-resistant Enterococcus faecium (VREF) method was utilized to estimate the total numbers of nuclei in the heart, as previously described. NV was an estimate of the numerical CM density, and VREF is the reference left ventricle volume. The total number of CMs was calculated based on the number of CM nuclei and the multinucleation level. The analysis was performed by confocal laser scanning microscopy (Carl Zeiss).

### Estimation of total number of CMs in vivo

As previously described,8 tissue pieces (1-2 mm diameter) from the left ventricle were sampled. CM nuclei were stained with antibodies against PCM-1, and nuclei were stained with DAPI. The cytoplasm of CMs was stained with cTnT. To facilitate the identification of the cell borders, WGA was added. Using the CAST software, the serial sections could be analyzed all together. In each block, the number of CMs was calculated, and serial sections were counted to estimate the total number of CMs. For each animal, 4 different tissue blocks were analyzed.

### Flow cytometry

After transfected with siRNAs or adenovirus, isolated CMs were cultured for 48h and then collected and fixed with cold 70% ethanol. Samples were centrifuged for 15 min at ×1200g. Cell pellets were re-suspended in FxCycle PI/RNase Staining Solution (Thermo Fisher Scientific) and analyzed on MoFlo XDP (Cell Sorter).

### Time-lapse videos

After the P7 CMs were transfected with Adv-Snhg1 or Adv-NC for 48 h, the CMs were labeled with tetramethylrhodamine ethyl ester (TMRE), a fluorescent dye that labels mitochondria. Then, the images were imaged for 12 h at 10 min intervals. Live-cell imaging was performed using a Delta Vision Elite system (Applied Precision) on an Olympus IX71 in verted microscope, running Soft WorX6.0. Time-lapse imaging was carried out for 12 h at 10 min intervals, and acquired at a 10x magnification (10/0.3 NA objective) with a Cool Snap HQ2CCD (charge-coupled device) camera (Roper Scientific).

For the P7 CMs isolated from the Myh6-mCherry transgenic mice, after transfection with Adv-Snhg1 for 48 h cells were imaged for 12-h at 10 min intervals. Live-cell imaging was performed using a Delta Vision Elite System (Applied Precision), on an Olympus IX71 in verted microscope, running Soft WorX6.0. Time-lapse imaging was carried out for 12 h at 10 min intervals, and acquired at 10 magnifications (10/0.3 NA objective) with a Cool Snap HQ2CCD (charge-coupled device) camera (Roper Scientific).

### RNA-seq analyses

RNA-seq and genome-wide transcriptome analyses. RNA was collected from the hearts of mice injected with AAV9-Snhg1 or AAV9-NC. RNA-seq experiments were performed by GENE DENOVE (Beijing, China). Briefly, the total RNA was isolated from fresh ventricular tissue using TRIzol (Invitrogen). The RefSeq and Ensembl transcript databases were used as the annotation references for mRNA analyses. The clustering of the index-coded samples was performed on a cBot Cluster Generation System using the TruSeq PE Cluster Kit v3-cBot-HS (Illumina) according to the manufacturer’s instructions. Analysis of differential expression was performed using the edgeR R package (3.12.1). The P values were adjusted using the Benjamini and Hochberg method. GO and KEGG pathway analyses were implemented using the cluster Profiler R package. The hierarchical clustering heat map was generated with the ggplot library.

### Pull-down assay

The probes of Snhg1 and its antisense RNA for the RNA pull-down assay were designed by Gzscbio Co., Ltd. (Guangzhou, China). The probes for the DNA pull-down assay were synthesized by Gzscbio Co., Ltd. Isolated CMs were washed in ice-cold PBS, lysed in 0.5 mL co-IP buffer, and incubated with 3 μg biotinylated DNA oligo probes against the Snhg1 back-splice sequence at room temperature for 4 h. Next, the CMs were incubated in 50 μL washed streptavidin-coated magnetic beads (Invitrogen) at room temperature for another hour. RNase-free BSA and yeast tRNA (Sigma) were used to prevent the nonspecific binding of RNA and protein complexes. RNA bound to beads was extracted with TRIzol, while the bound protein was analyzed by Western blotting. The specific bands were extracted and then analyzed by mass spectrometry or Western blotting.

### RIP

A biotin-labeled anti-PTEN antibody (1:1000, #4370, cell signaling technology,) and IgG antibodies were added to cell extracts and incubated overnight at 4°C. Streptavidin-coated magnetic beads were then added and incubated for 4 h at 4°C. Magnetic beads were pelleted, washed, and resuspended in 1 mL TRIzol. Isolated RNA was reverse transcribed to cDNA and then analyzed by RT-qPCR. The PCR cycling parameters were as follows: an initial denaturation of 95°C for 5 min followed by 40 cycles of 94°C for 10 s and 65°C for 40 s, with a final extension at 72°C for 10 min.

### Echocardiography

To evaluate cardiac function and dimensions, transthoracic two-dimensional echocardiography was performed on mice anesthetized with 2% isoflurane using a Vevo 2100 Imaging System (Visual Sonics, Ontario, Canada) equipped with a 40-MHz probe. M-mode tracings in the parasternal short axis view were used to measure the left ventricular internal diameter at end-diastole (LVEDd) and end-systole (LVESd), which were used to calculate left ventricular fractional shortening (LVFS) and the left ventricular ejection fraction (LVEF).

### Masson’s trichrome staining

Formalin-fixed, paraffin-embedded heart tissue slides were deparaffinized via xylene and rehydrated through sequential incubations in ethanol (100%, 100%, 90%, 80%, and 70%) and water. The slices were incubated in Weigert hematoxylin iron for 5 min, differentiated in hydrochloric acid (HCl)-ethanol, incubated in ponceau acid fuchs for 5 min, phosphomolybdic acid for 5 min, and aniline blue or light green (Leagene, Beijing, China) for 5 min. The fibrotic area was measured as the percentage of the total left ventricular area showing fibrosis and quantified with ImageJ software (NIH, Bethesda, MD, USA).

### Immunohistochemistry

The MI heart sections were incubated with antibodies against CD31 (Abcam, Cambridge, UK) at 4°C overnight, and then with secondary antibodies at 4°C for 1 h and detected with 3, 3′-diaminobenzidine. The sections were counterstained with hematoxylin. Immunohistochemistry images were captured with an Olympus BX51 microscope (Tokyo, Japan).

### RNA isolation and RT-qPCR

Total RNA from isolated CMs or dissected ventricular heart tissue samples was extracted using the E.Z.N.A. ® Total RNA Kit II (Omega Biotek, Norcross, GA, USA) according to the manufacturer’s instructions. Cytoplasmic and nuclear RNAs were separated with an RNeasy Midi Kit (Qiagen, Hilden, Germany). The RNAs were treated with DNase I (Invitrogen) to prevent DNA contamination. cDNA was reverse-transcribed from 1 μg of total RNA using the PrimeScript™ RT Master Mix (TaKaRa, Shiga, Japan). For reverse transcription of Snhg1, strand-specific primers and a reverse primer of Snhg1 were used as reverse transcription primers. RT-qPCR was performed with SYBRs Premix Ex Taq™ Kit (TaKaRa) on a Lightcycler 480 (Roche, Basel, Switzerland). Briefly, the 20μl reaction mixtures were incubated at 95°C for 30s for the initial 3 denaturation, followed by 40 cycles at 95°C for 5s and 60°C for 34s. β-actin was used as a housekeeping control gene to normalize gene expression using the ^ΔΔ^Ct method. All primers were designed by Sangon Biotech (Shanghai, China). Primer sequences are shown in Table S1.

### Western blotting

Isolated mouse CMs or dissected mouse ventricular heart tissue samples were lysed in ice-cold radio immunoprecipitation assay buffer (Ding guo Chang sheng, Beijing, China) with protease inhibitors and phosphatase inhibitors. Protein concentrations were determined with the BCA Protein Quantitative Analysis Kit (Fudebio-tech, Hangzhou, China). Standard Western blotting were performed.^10^ Briefly, protein samples were separated by 8-12% sodium dodecyl sulfate-polyacrylamide gel electrophoresis (SDS-PAGE) and transferred onto polyvinylidenedifluoride membranes (Millipore, Billerica, MA, USA). The membranes were incubated at room temperature for 2 h in blocking buffer (5% BSA in Tris-buffered saline and Tween 20 [TBST] buffer). After blocking, the membranes were incubated with primary antibodies overnight at 4°C. Primary antibodies against the following proteins were used: PTEN (Cell Signaling Technology, #9188), p-Akt (Cell Signaling Technology, #4060), Akt (Cell Signaling Technology, #9272), pGsk3β (Abcam, ab131097), c-Myc (Cell Signaling Technology, #18583,Proteintech, 10828-1-AP), and β-actin (Biosynthesis, bs-0061R). After the membranes were washed three times with TBST, they were incubated with donkey anti-rabbit IgG (Abcam, ab16284) for 1 h at room temperature. The membranes were developed with the electrochemiluminescence method according to the manufacturer’s instructions (Millipore) and detected on a chemiluminescence imaging GeneGnome XRQ System (Syngene, Bangalore, India). To calculate the relative density, ImageJ software was used, and the intensity of each band was normalized to that of β-actin.

### Luciferase assay

Luciferase assays were conducted with the Dual-luciferase Reporter Kit (Promega). The pRL vector constitutively expressing Renilla luciferase was used to normalize for transfection efficiency. A total of 2 × 10^5^ CMs were plated in 12-well dishes 24 h before transfections. On the day of transfection, each well was transfected with pRL, pGL3 Basic (to assess basal reporter activity) or Myc promoter-pGL3 and the indicated plasmids. Twenty-four hours later, luciferase activity was measured using the Wallac 1450 MicroBeta TriLux System (Perkin Elmer). Experiments were carried out 3 times in triplicates, and error bars represent SD.

---

**Figure S1.**
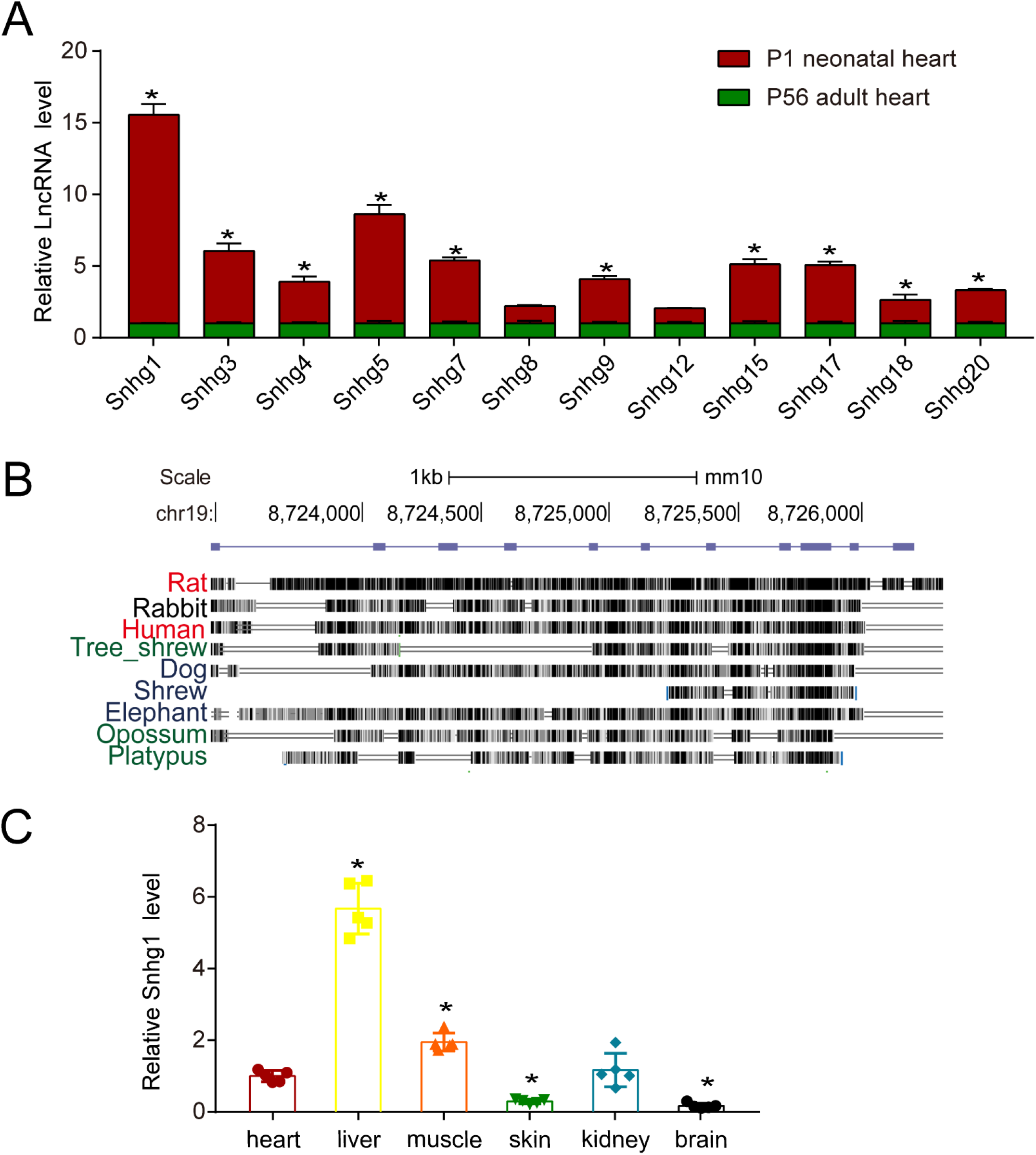
Snhg1 is highly expressed in neonatal hearts and conserved across mammalians. (A) RT-qPCR analysis of Snhg1, 3, 4, 5, 7, 8, 9, 12, 15, 17, 18, and 20 in P1 and adult mouse hearts as detected by RT-qPCR, n=5. (B) Conservation analysis of Snhg1. (C) RT-qPCR analysis of Snhg1 levels in P1 mouse heart, liver, muscle, skin, kidney, and brain tissues, n=5. All data are expressed as the mean ± SD, \**P*<0.05 using t-tests in A, one-way ANOVA test followed by LSD post hoc test in C.

**Figure S2.**
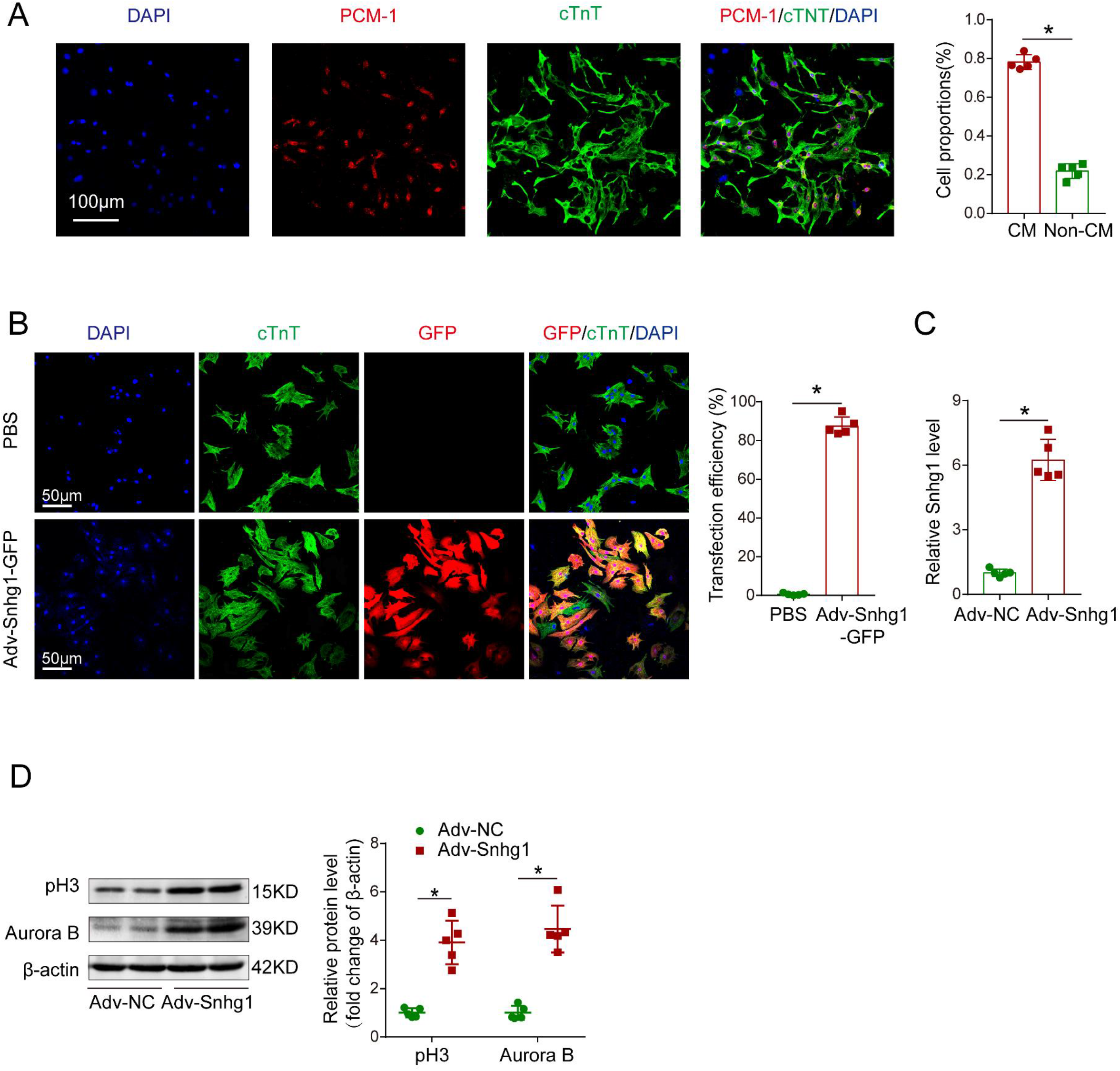
Adenovirus-mediated Snhg1 overexpression promotes P7 CM proliferation *in vitro*. (A) The purity of isolated P7 CMs is approximately 80%, and isolated CMs were stained with cTnT, PCM-1, and DAPI, n=5. (B) GFP/cTnT double-staining in P7 CMs after transfection with PBS or Adv-Snhg1-GFP, n=5. (C) Quantification of Snhg1 expression by RT-qPCR in isolated P7 CMs transfected with Adv-NC or Adv-Snhg1, n=5. (D) Western blotting analysis and quantification of pH3 and Aurora B protein levels in isolated P7 CMs transfected with Adv-NC or Adv-Snhg1. β-actin was used as a loading control, n=5. All data are expressed as the mean ± SD, \**P*<0.05 using t-tests in A-D.

**Figure S3.**
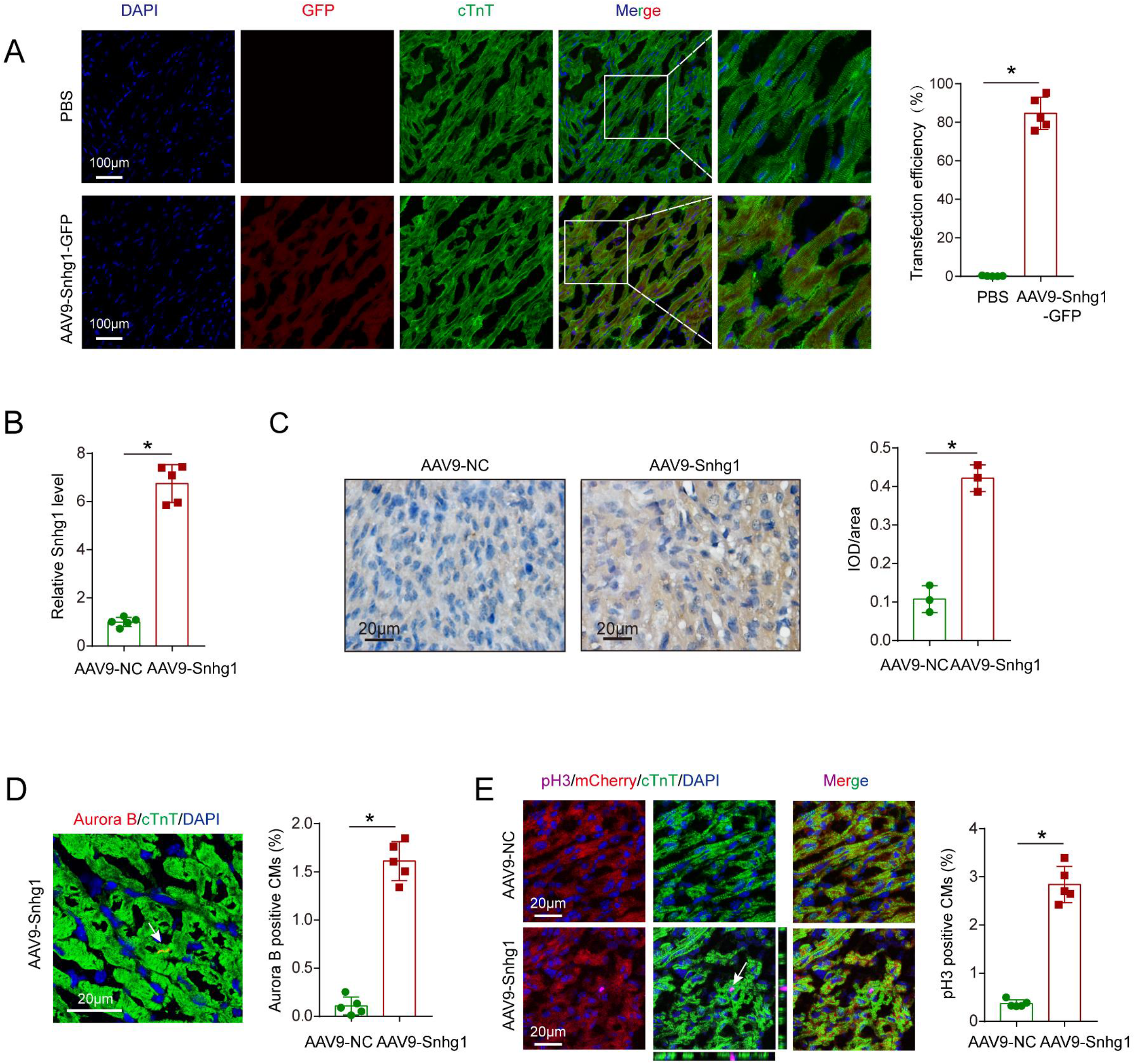
Snhg1 overexpression promotes P7 CM proliferation *in vivo*. (A) The transfection efficiency of AAV9-Snhg1 was approximately 80%, as revealed by GFP and CM specific marker staining, n=5. (B) Quantification of Snhg1 expression by RT-qPCR in P7 mouse hearts injected with AAV9-NC or AAV9-Snhg1, n=5. (C) ISH assays confirmed that Snhg1 was significantly increased in P7 mouse hearts injected with AAV9-Snhg1 or AAV9-NC. the brown dot cluster indicates Snhg1, n=3. (D) Immunostaining and quantification of Aurora B in P7 mouse hearts injected with AAV9-NC or AAV9-Snhg1, n=5. (E) Immunostaining and quantification of pH3-positive CMs in P7 Myh6-mCherry transgenic mice injected with AAV9-NC or AAV9-Snhg1, n=5. All data are expressed as the mean ± SD, \**P*<0.05 using t-tests in A-E.

**Figure S4.**
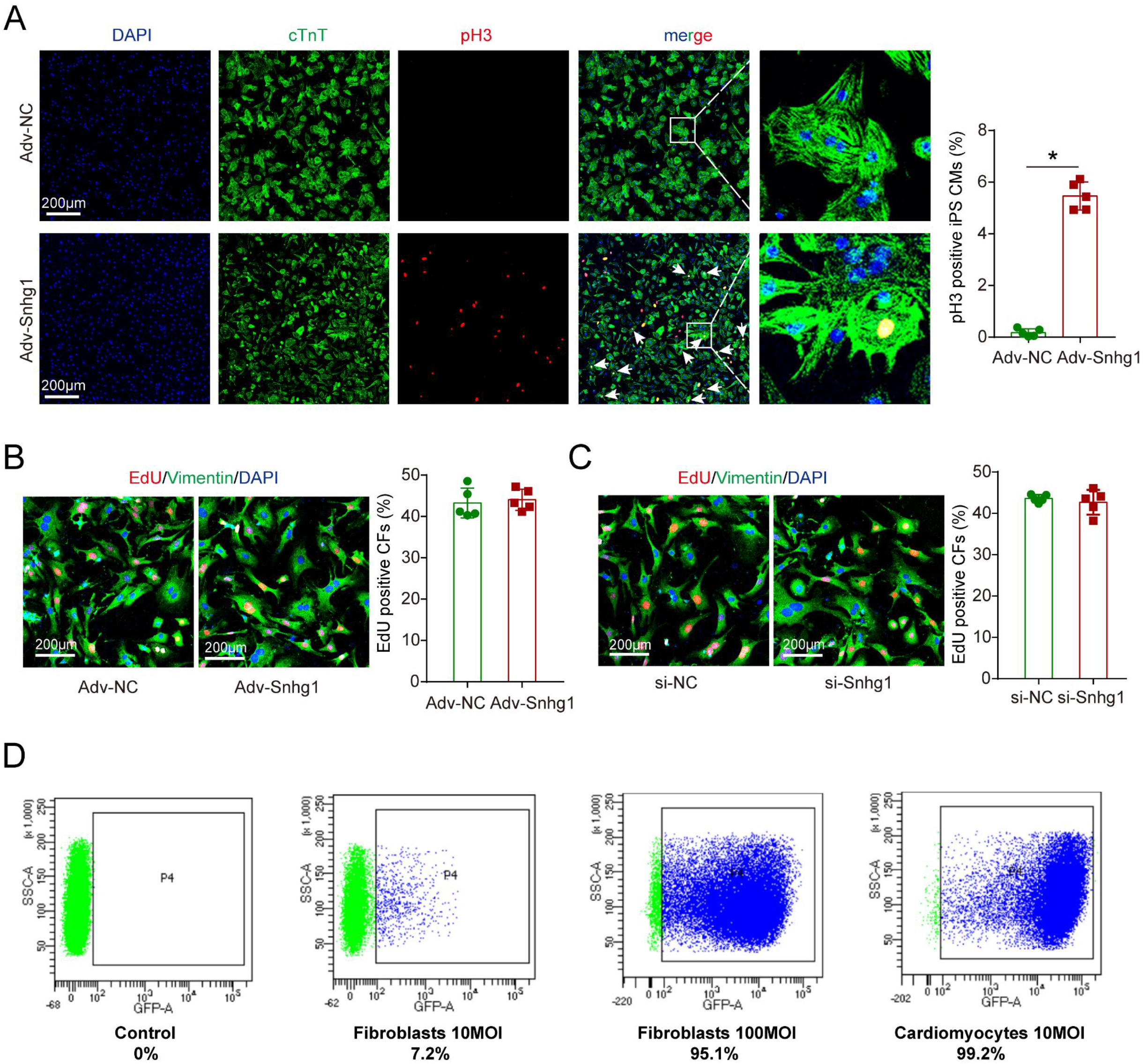
Snhg1 promotes human iPS-derived CM proliferation but not CF proliferation. (A) Immunocytochemistry of 60-day-old human iPS-derived CMs infected with Adv-NC or Adv-Snhg1 and immunostained 48 h later with antibodies against pH3, cardiac troponin T, and DAPI to mark nuclei, n=5. (B) EdU immunofluorescence staining in CFs transfected with Adv-NC or Adv-Snhg1 and quantification of EdU-positive CFs, n=5. (C) EdU immunofluorescence staining in CFs transfected with si-NC or si-Snhg1 and quantification of EdU-positive CFs, n=5. (D) Representative FACS plots showing infection efficiency of GFP adenovirus in Thy1+ cardiac fibroblasts infected with 10 or 100 MOI, compared to CMs infected with 10 MOI of the adenovirus. All data are expressed as the mean ± SD, \**P*<0.05 using t-tests in A-C.

**Figure S5.**
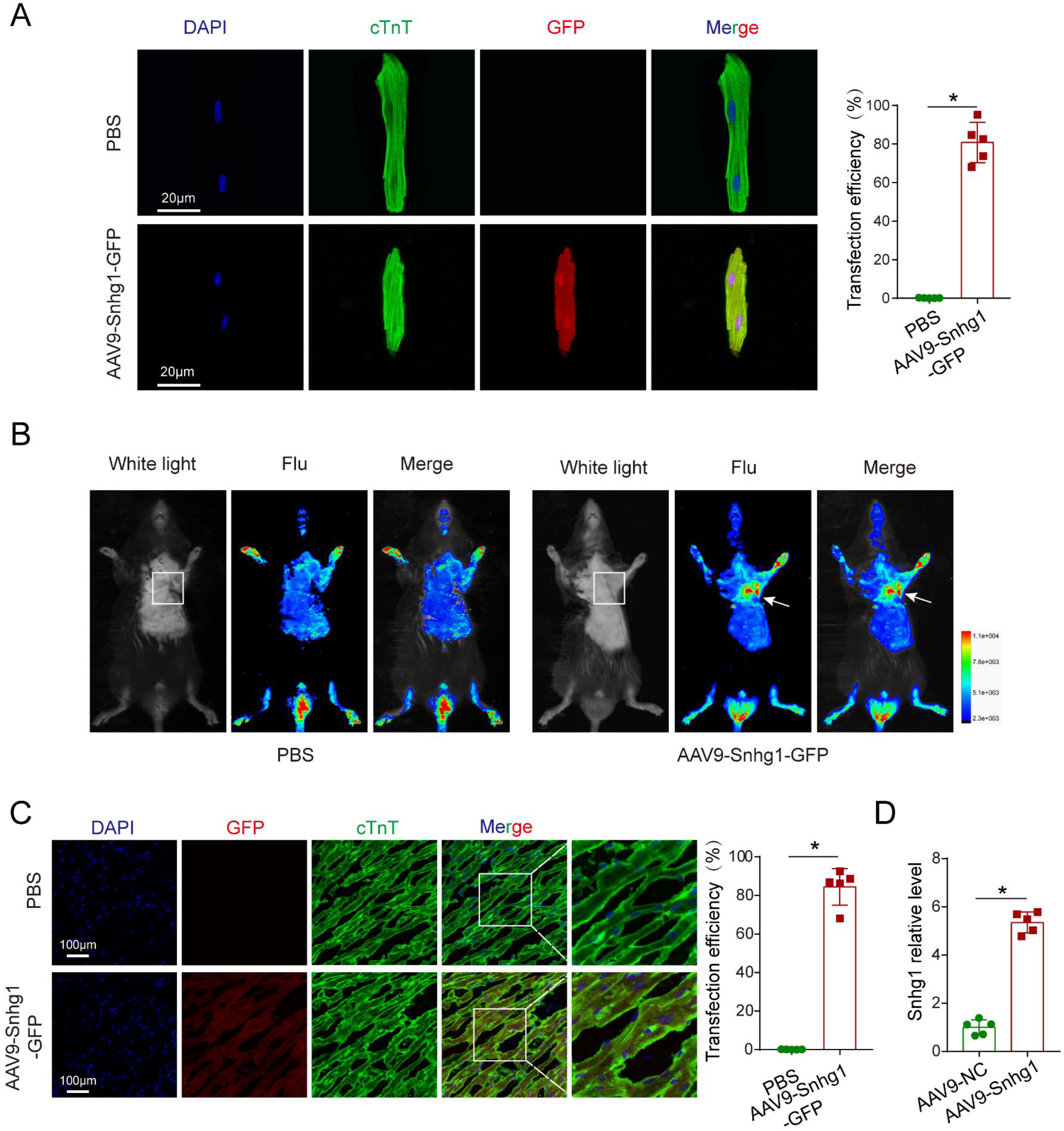
The transfection efficiency of AAV9-Snhg1. (A) The transfection efficiency of isolated adult CMs in AAV-Snhg1 injected mouse was approximately 80% as revealed by GFP and CM specific marker staining, n=5. (B) Representative In vivo bioluminescent images and Bright field captured on day 14 after injection with GFP-labelled AAV9-Snhg1 virus. Square indicates thoracic incision, arrow indicates the heart with GFP fluorescence. (C) GFP/cTnT double-staining in mouse adult hearts injected with PBS or AAV9-Snhg1-GFP, n=5. (D) Quantification of Snhg1 expression by RT-qPCR in adult mouse hearts injected with AAV9-NC or AAV9-Snhg1, n=5. All data are expressed as the mean ± SD, \**P*<0.05 using t-tests in A and C-D.

**Figure S6.**
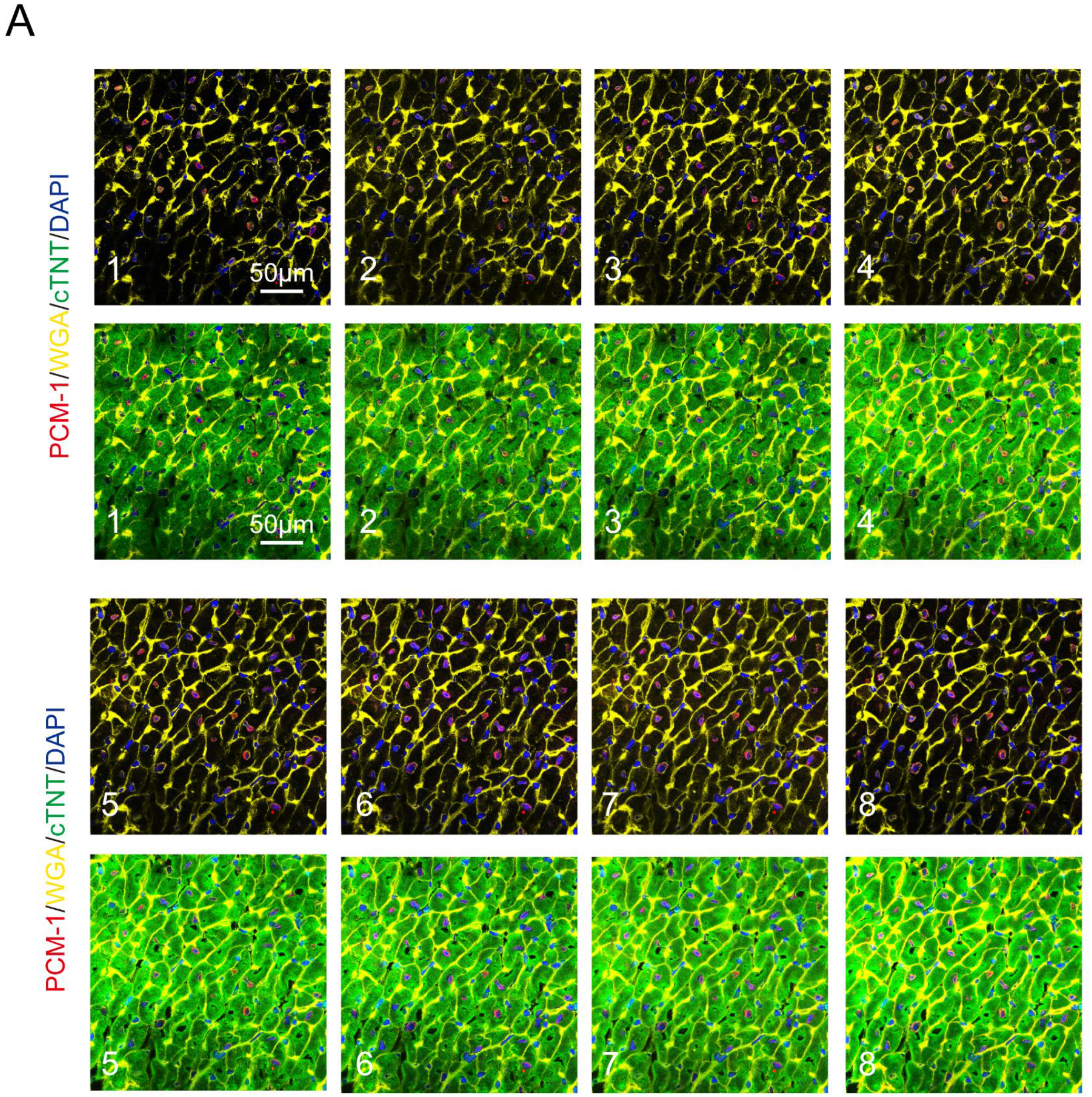
Snhg1 induces adult CM proliferation. (A) Stereological analysis revealed the number of CMs in adult mice injected with AAV9-NC or AAV9-Snhg1. Sections 1-8 out of 8 serial sections are shown, n=5.

**Figure S7.**
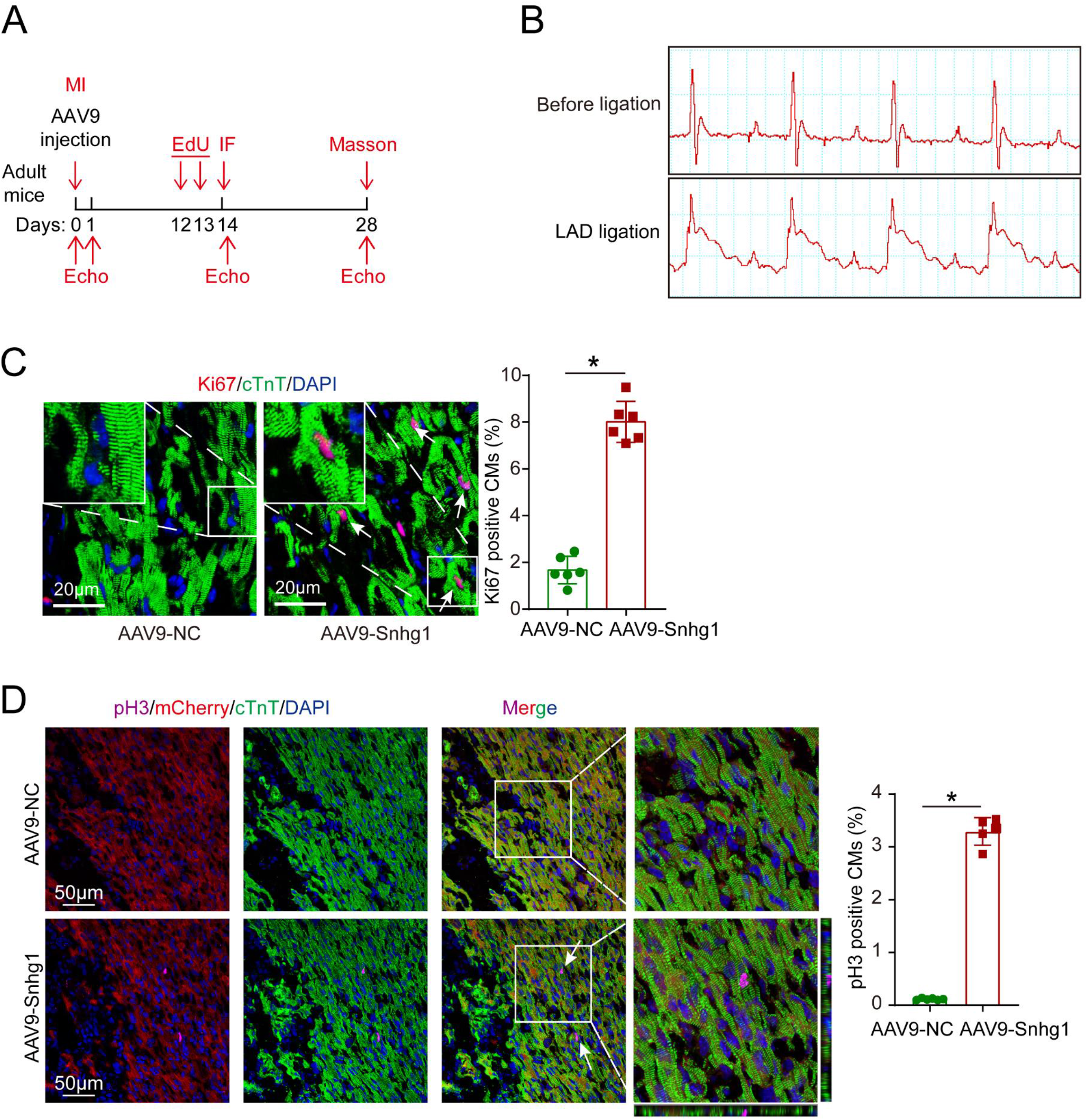
Snhg1 improves adult cardiac function post-MI. (A) Schematic of the MI experiments in adult mouse hearts injected with AAV9. Echo, echocardiography. (B) The adult mouse MI model was confirmed using an electrocardiogram ST-segment elevation. (C) Immunofluorescence of Ki67 positive CMs in AAV9-NC and AAV9-Snhg1-injected adult mouse hearts at 14 days after MI, n=6. (D) Immunostaining and quantification of pH3-positive CMs in P7 Myh6-mCherry transgenic mice injected with AAV9-NC or AAV9-Snhg1, n=5. All data are expressed as the mean ± SD, \**P*<0.05 using t-tests in C-D.

**Figure S8.**
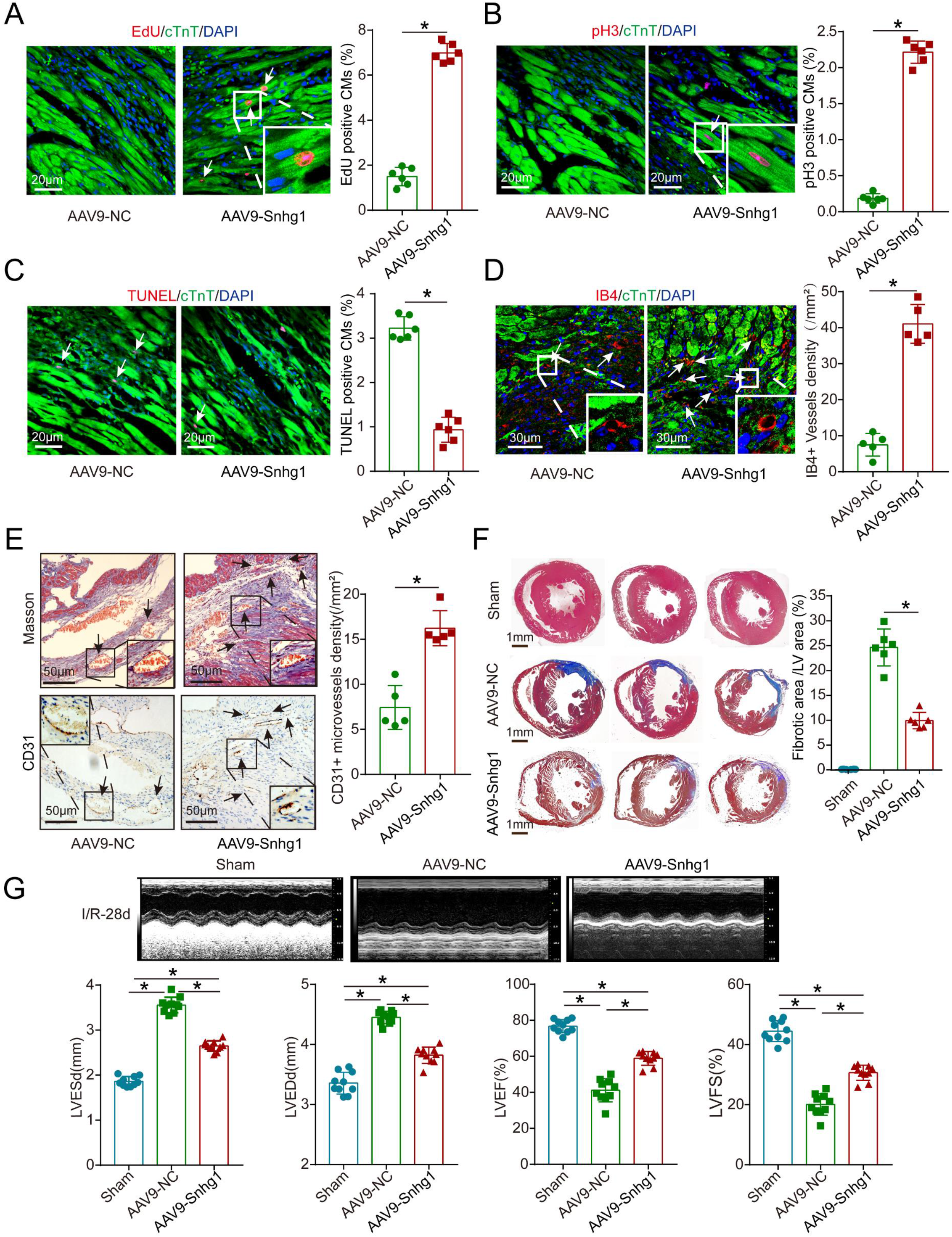
Snhg1 improves adult cardiac function in the I/R mouse model. (A-B) Immunofluorescence and quantification of EdU and pH3 for AAV9-NC and AAV9-Snhg1-injected adult mouse hearts 14 days after I/R, positive CMs are indicated by arrows, n=6. (C) Immunofluorescence staining and quantification of TUNEL-positive CMs in the AAV9-NC and AAV9-Snhg1-injected adult mouse hearts 14 days after I/R. TUNEL-positive CMs are indicated by arrows, n=6. (D) Immunofluorescence and quantification of IB4-positive cells for AAV9-NC and AAV9-Snhg1-injected adult mouse hearts 14 days after I/R. IB4-positive cells are indicated by arrows, n=5. (E) Representative images of Masson’s trichrome staining and immunohistochemistry for CD31 in AAV9-NC or AAV9-Snhg1-injected adult mice 28 days after I/R and quantification of CD31-positive cells, n=5. (F) Representative images of Masson’s trichrome-stained heart sections in Sham or AAV9-NC or AAV9-Snhg1-injected adult mice 28 days after I/R and quantification of infarct size, n=6. (G) Representative images of echocardiography analysis on Sham or AAV9-NC or AAV9-Snhg1-injected adult mouse hearts at 28 days after I/R and quantification of LVESd, LVEDd, LVEF, and LVFS, n=10. All data are expressed as the mean ± SD, \**P*<0.05 using t-tests in A-E, one-way ANOVA test followed by LSD post hoc test in F-G.

**Figure S9.**
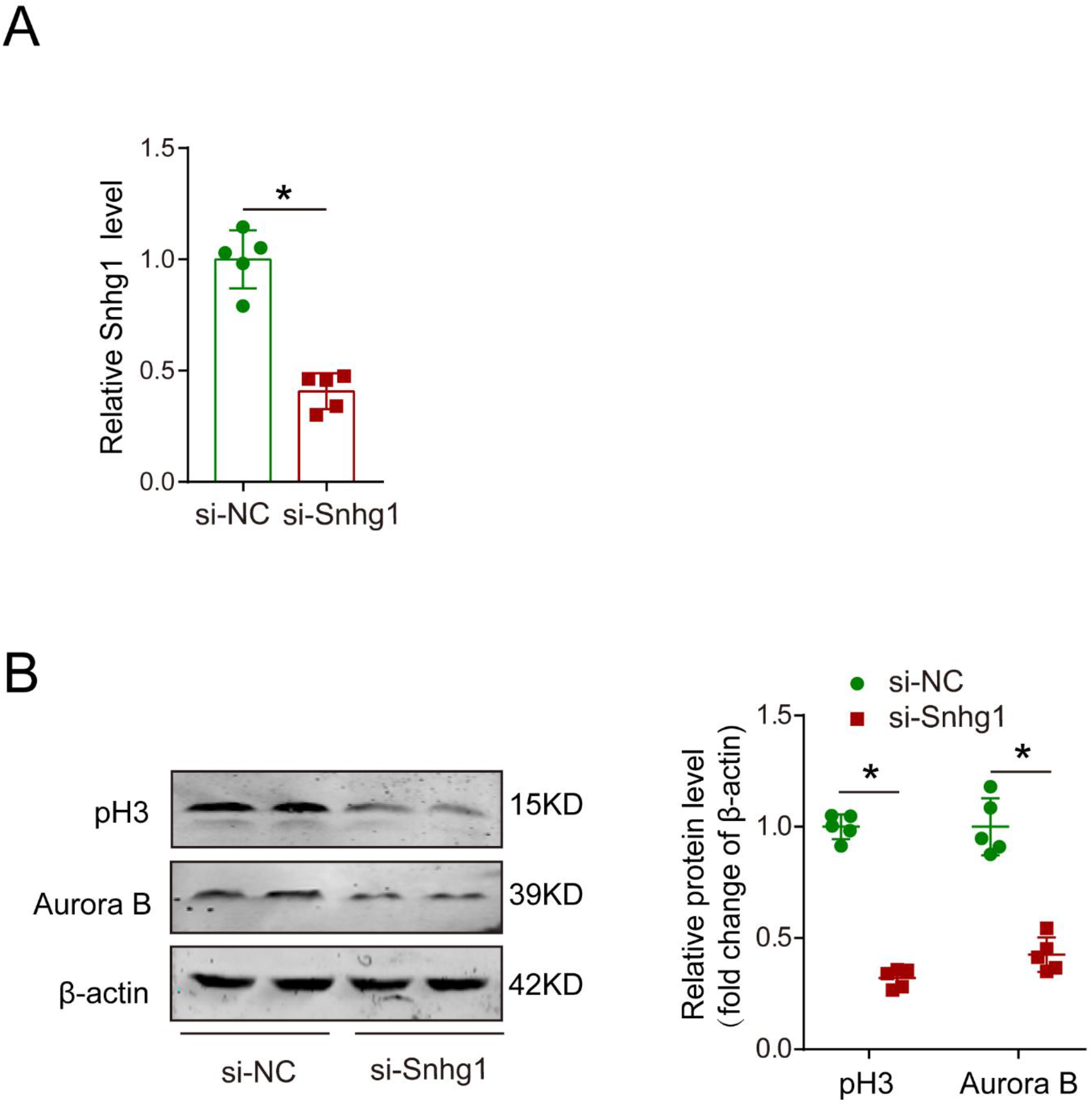
Snhg1 depletion decreased neonatal CM proliferation *in vitro*. (A) RT-qPCR of Snhg1 in isolated P1 CMs transfected with si-NC or si-Snhg1, n=5. (B) Western blotting analysis of pH3 and Aurora B protein levels in isolated P1 CMs transfected with si-NC or si-Snhg1, n=5. All data are expressed as the mean ± SD, \**P* <0.05 using t-tests in A-B.

**Figure S10.**
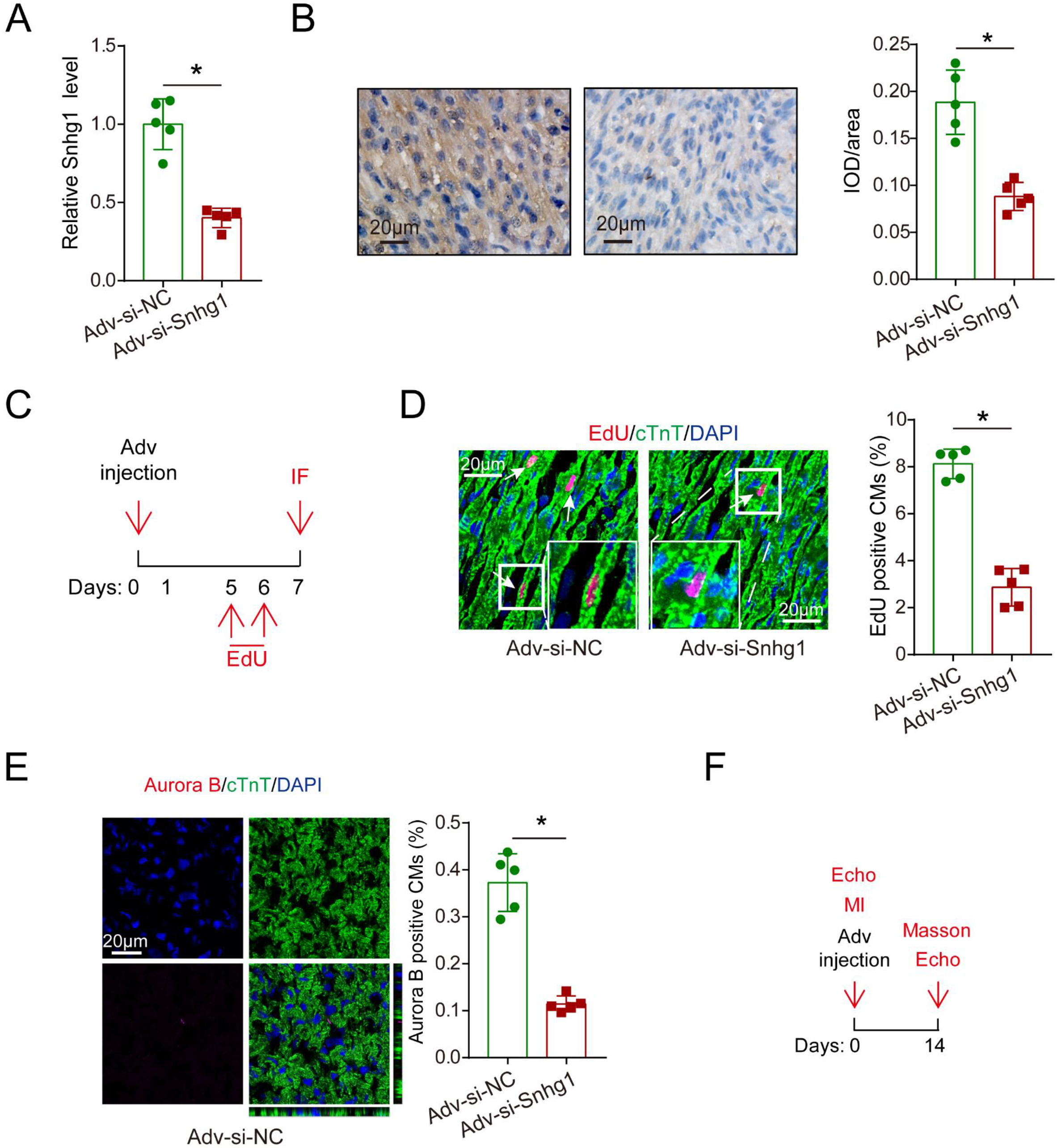
Snhg1 depletion decreased neonatal CM proliferation *in vivo*. (A) RT-qPCR of Snhg1 expression in neonatal mouse hearts injected with Adv-si-NC or Adv-si-Snhg1, n=5. (B) *ISH* assays of Snhg1 expression in neonatal mouse hearts injected with Adv-si-NC or Adv-si-Snhg1. n=5. (C) Schematic of experiments in neonatal mouse hearts injected with adenovirus. (D) Immunofluorescence of EdU in neonatal hearts injected with Adv-si-NC or Adv-si-Snhg1, n=5. (E) Immunofluorescence and quantification of Aurora B-positive CMs in neonatal hearts injected with Adv-si-NC or Adv-si-Snhg1, n=5. (F) Schematic of MI experiments in neonatal mouse hearts injected with adenovirus. Echo, echocardiography. All data are expressed as the mean ± SD, \**P*< 0.05 using t-tests in A-B and D-E.

**Figure S11.**
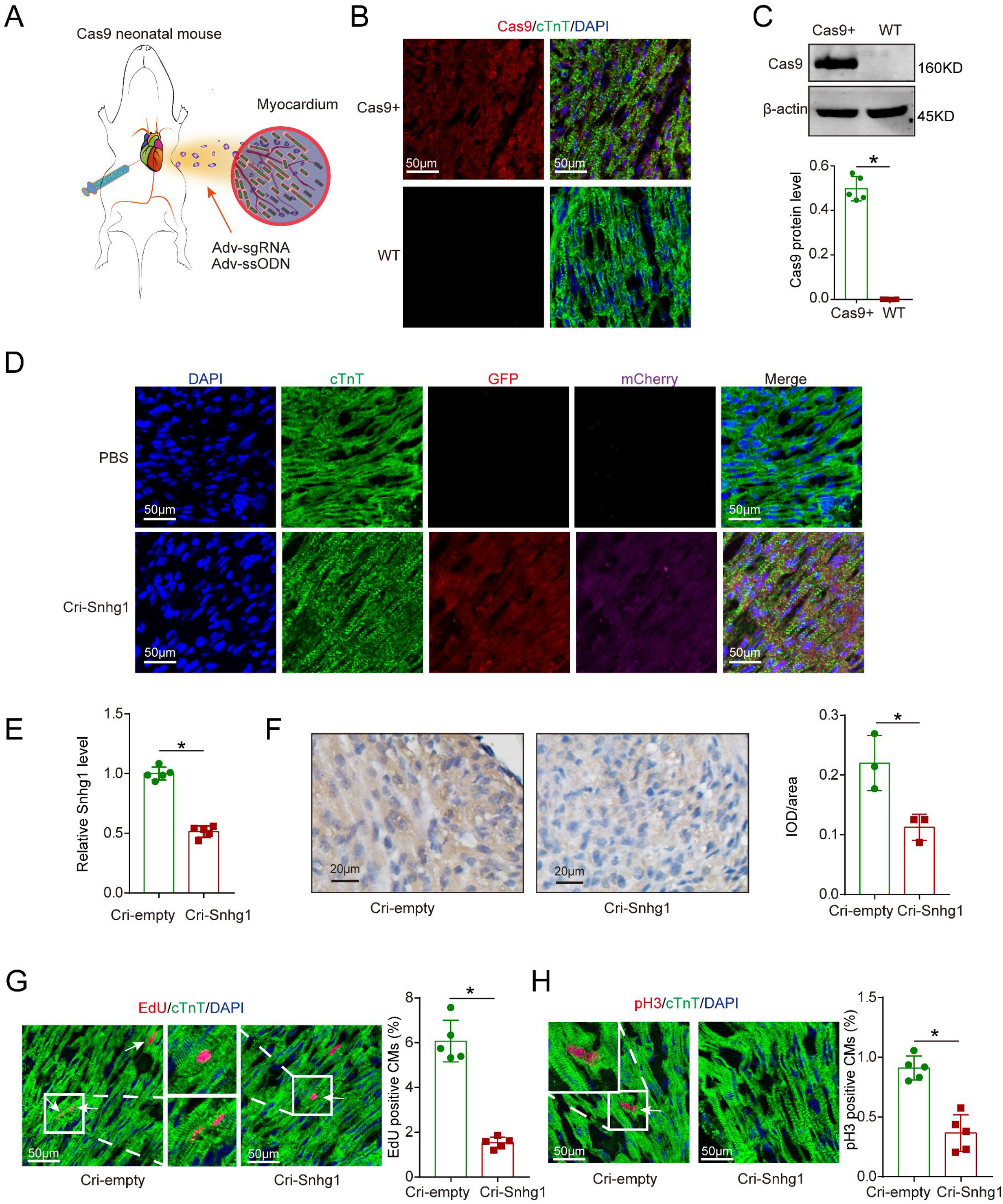
Generation of Snhg1-deficient mice using CRISPR-Cas9 technology. (A) Schematic illustrating the procedure for the delivery of Adv expressing sgRNA and ssODN into the myocardium tissue of Cas9 knock in mice. (B) Immunostaining of Cas9 in Cas9 transgenic mouse hearts. Cas9+: Cas9 transgenic mice; WT: wild-type C57BL/6 mice. (C) Western blotting detected the Cas9 protein in Cas9 mouse hearts. Cas9+: Cas9 transgenic mice; WT: wild-type C57BL/6 mice, n=5. (D) Immunostaining of GFP in Cas9 mouse hearts at 7 days after injection with Adv -sgRNA(Snhg1)-GFP and Adv-ssODN(Snhg1)-mCherry. (E) RT-qPCR assays detecting Snhg1 expression in Cas9 mouse hearts at 7 days after injection with Adv -sgRNA(Snhg1)-GFP and Adv -ssODN(Snhg1)-mCherry(Cri-Snhg1). Cas9 mice injected with Adv -sgNC were used as the control group (Cri-empty). (F) ISH assays detecting Snhg1 expression in Cas9 mouse hearts 7 days after injection with Adv-sgRNA(Snhg1)-GFP and Adv -ssODN(Snhg1)-mCherry. Cas9 mice injected with Adv-sgNC were used as the control group (Cri-empty), n=3. (G-H) Immunofluorescence and quantification of EdU- and pH3-positive CMs for Adv-sgRNA(Snhg1)-GFP and Adv -ssODN(Snhg1)-mCherry-injected neonatal mouse hearts. Cas9 mice injected with Adv -sgNC were used as the control group (Cri-empty), n=5. All data are expressed as the mean ± SD, \**P*< 0.05 using t-tests in C and E-H.

**Figure S12.**
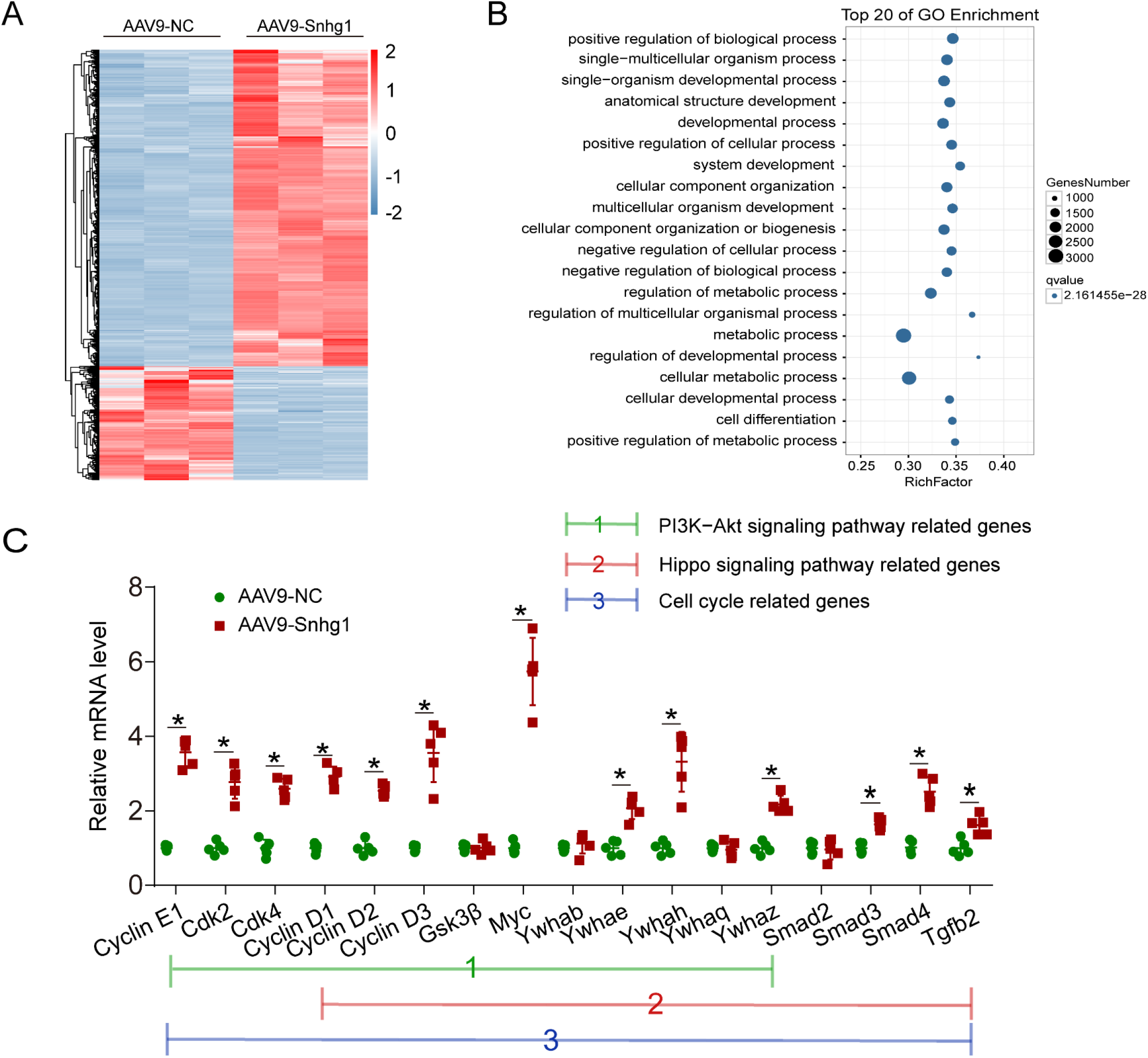
Next-generation RNA sequencing (RNA-seq) of Snhg1 overexpression and control group from P7 mice. (A) Hierarchical clustering of differentially expressed genes between AAV9-Snhg1 and control mimic-injected P7 hearts. Red and blue colors indicate up-regulated or down-regulated genes. (B) Scatter plot showing top 20 enriched GO terms of biological process for differentially expressed genes between AAV9-Snhg1 and control mimic-injected P7 hearts. (C) RT-qPCR analysis of the expression of differentially expressed genes related to the cell cycle, PI3K-Akt, and Hippo pathways, n=5. All data are expressed as the mean ± SD, \**P*< 0.05 using t-tests in C.

**Figure S13.**
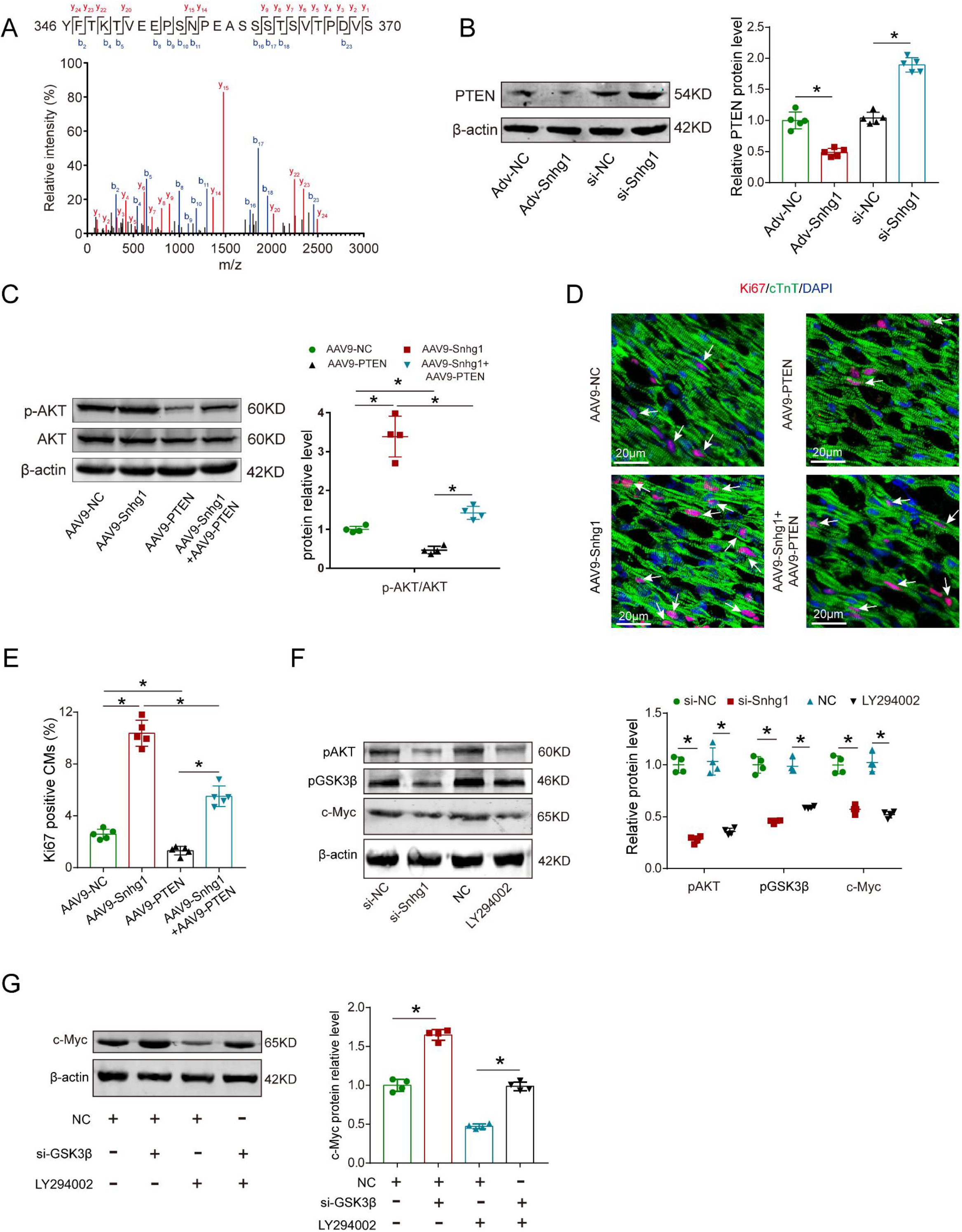
Snhg1 regulates CM proliferation through the PTEN/PI3K-Akt/c-Myc pathway. (A) NanoLC-MS/MS spectrum of PTEN peptides. (B) Western blotting analysis and quantification of PTEN protein levels in isolated CMs transfected with Adv-NC, Adv-Snhg1, si-NC, or si-Snhg1. β-Actin was used as a loading control, n=5. (C) Western blotting analysis and quantification of PI3K, p-Akt, and Akt protein levels in P7 mice injected with AAV9-NC, AAV9-Snhg1, AAV9-PTEN, and AAV9-Snhg1+AAV9-PTEN. β-Actin was used as a loading control, n=4. (D-E) Immunofluorescence and quantification of Ki67-positive CMs in P7 mice injected with AAV9-NC, AAV9-Snhg1, AAV9-PTEN, or AAV9-Snhg1+AAV9-PTEN, n=5. (F) Western blotting comparing p-AKT, p-GSK3β, and c-Myc levels in P7 CMs treated with si-NC, si-Snhg1, NC or LY294002. β-Actin was used as a loading control, n=4. (G) Western blotting comparing c-Myc levels in P7 CMs treated with NC, si-GSK3β, LY294002, or LY294002+ si-GSK3β. β-Actin was used as a loading control, n=4. All data are expressed as the mean ± SD, \**P*< 0.05 using t-tests in B, one-way ANOVA test followed by LSD post hoc test in C and E-G.

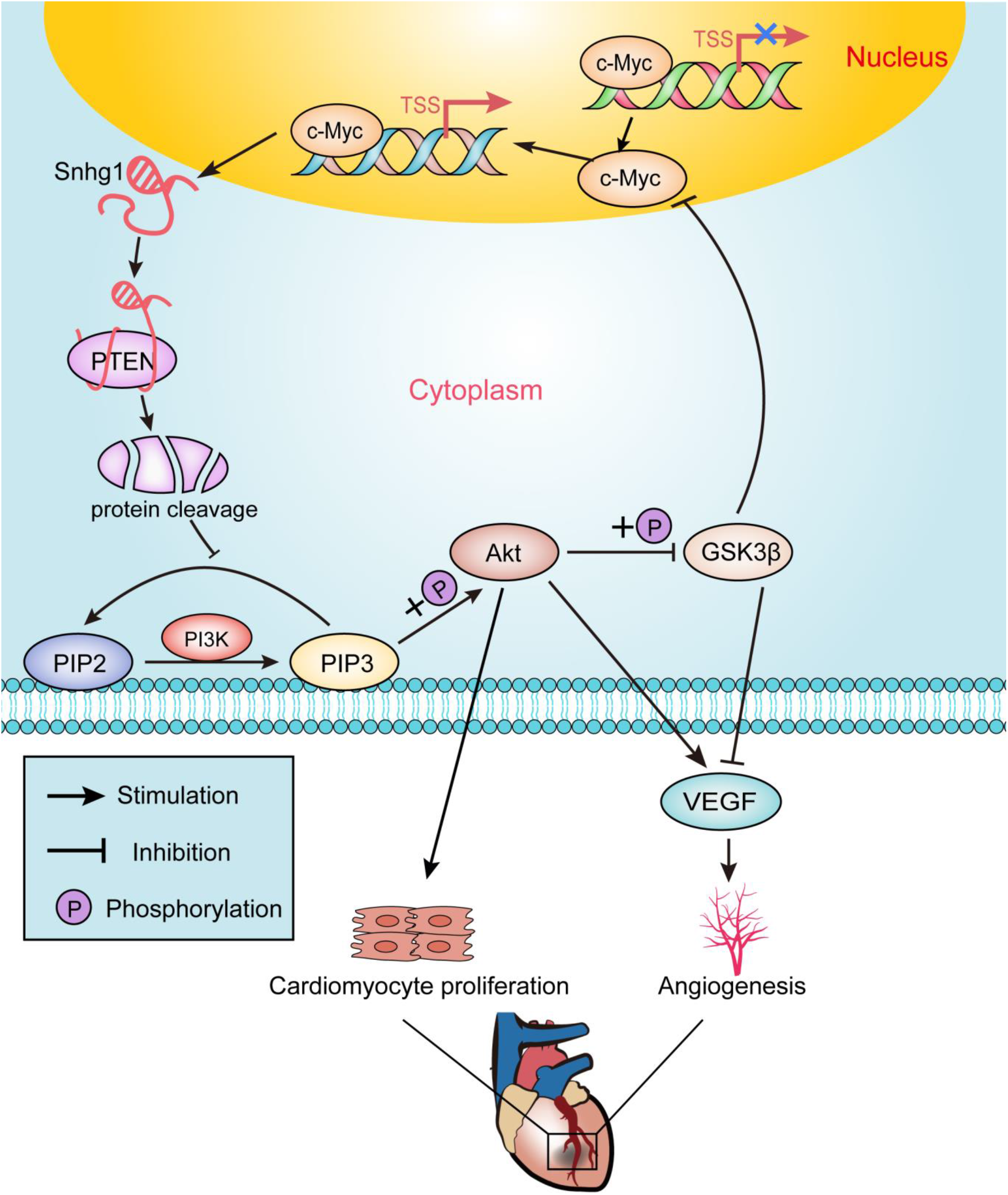

**Supplementary Table 1:**
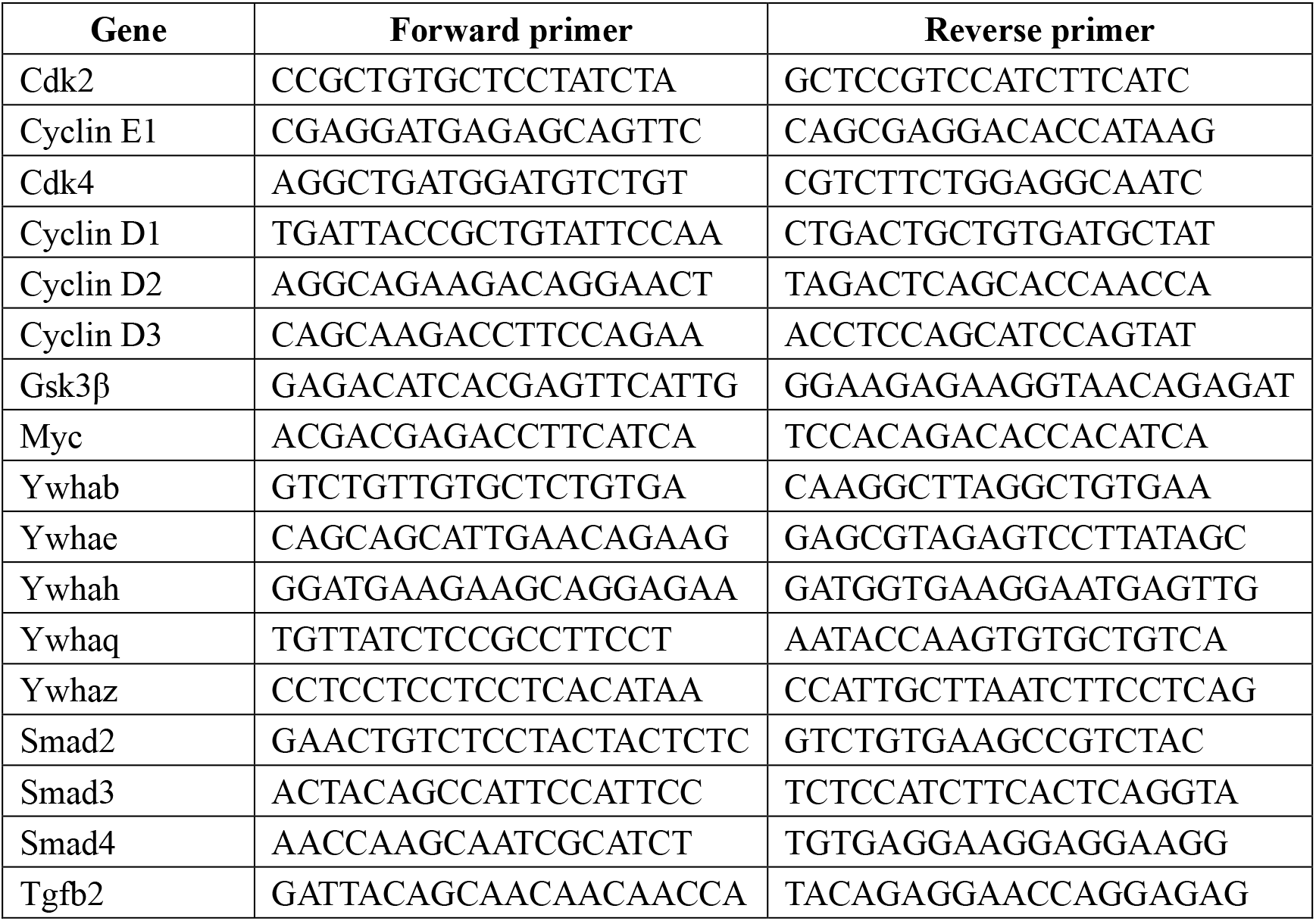
Sequences of primers for RT-qPCR

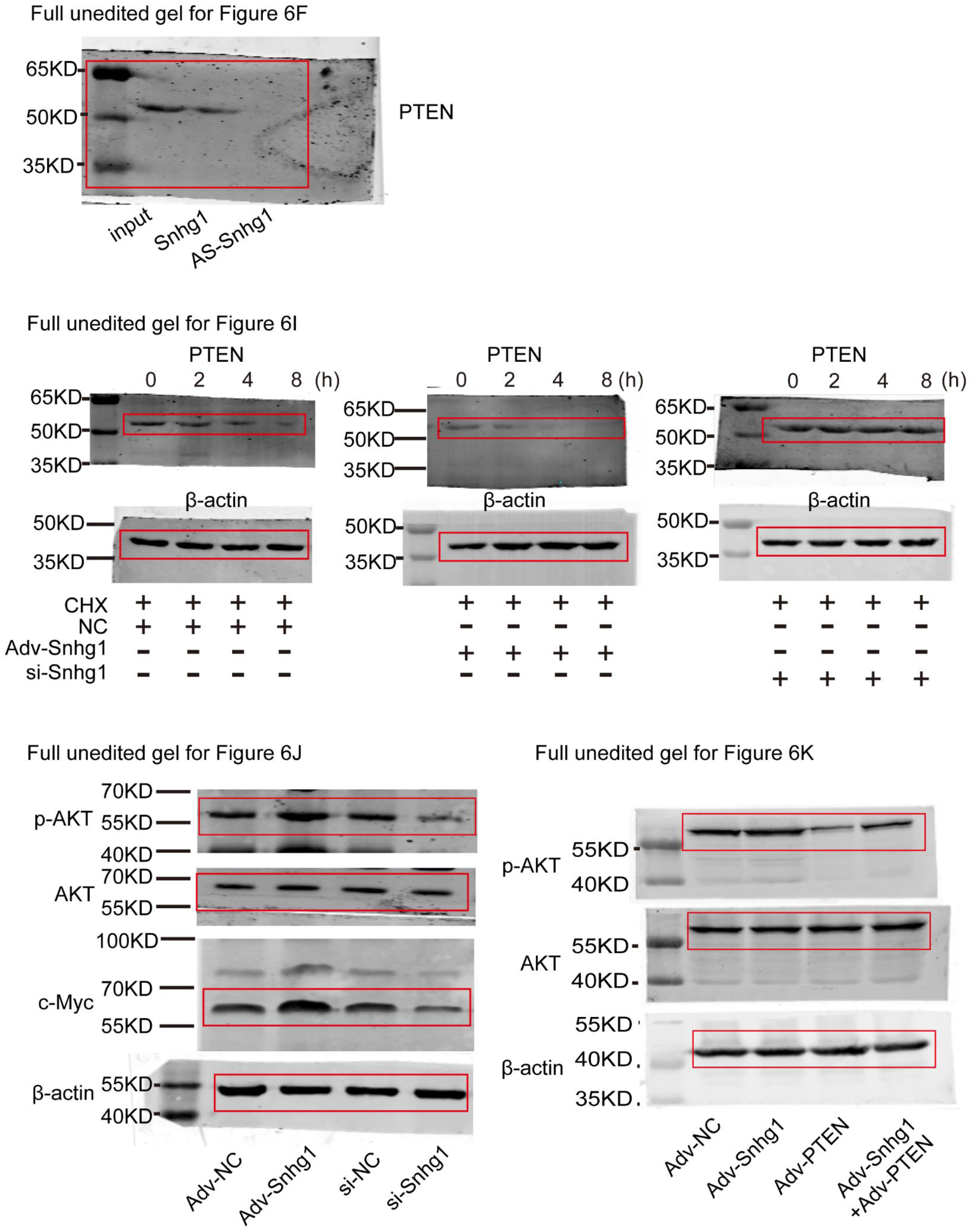
Entire unedited gel for representative cropped gels of Fig.6F,6I,6J,6K. Representative cropped gels are marked by red box.

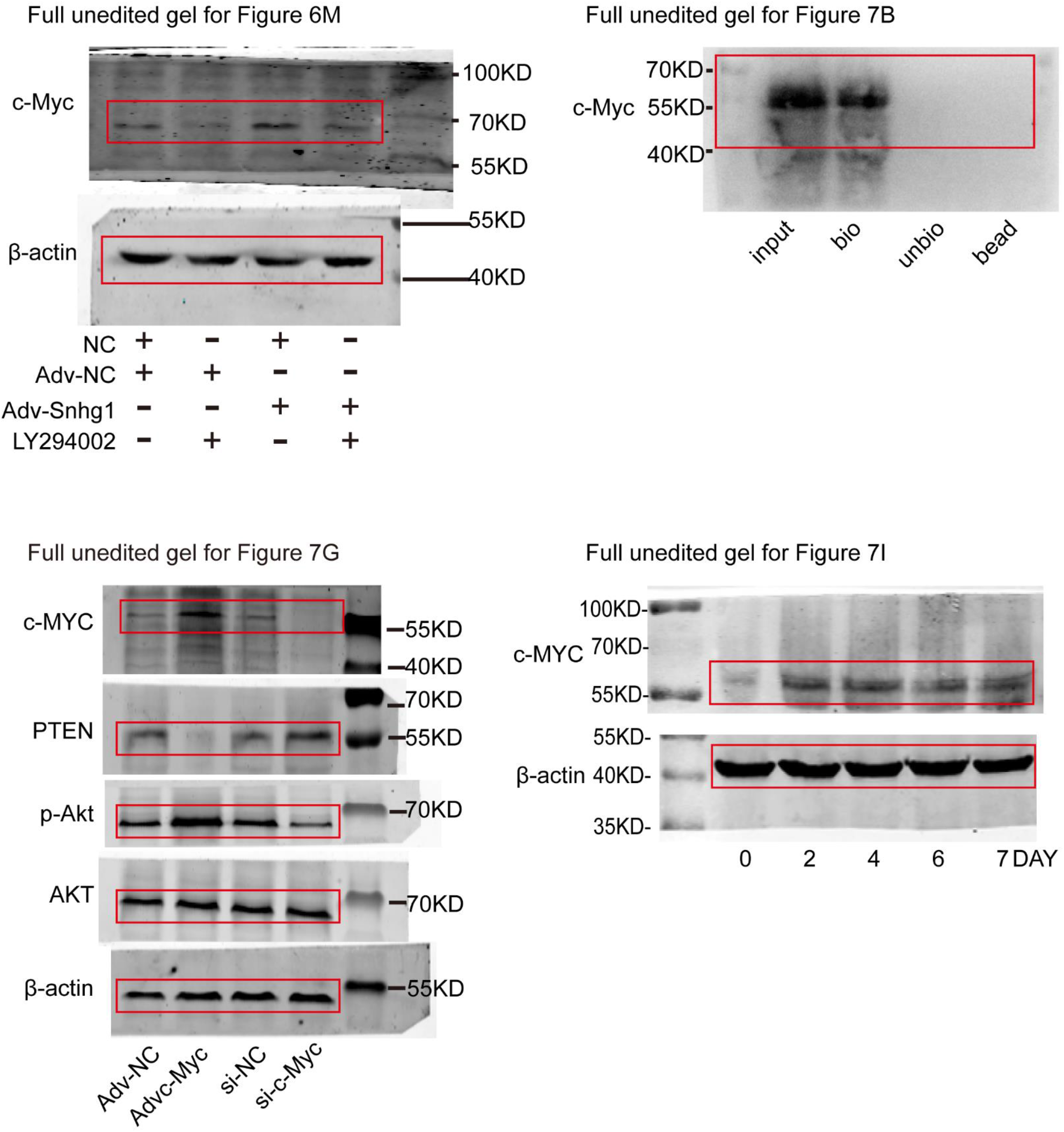
Entire unedited gel for representative cropped gels of Fig.6M,7B,7G,7I. Representative cropped gels are marked by red box.

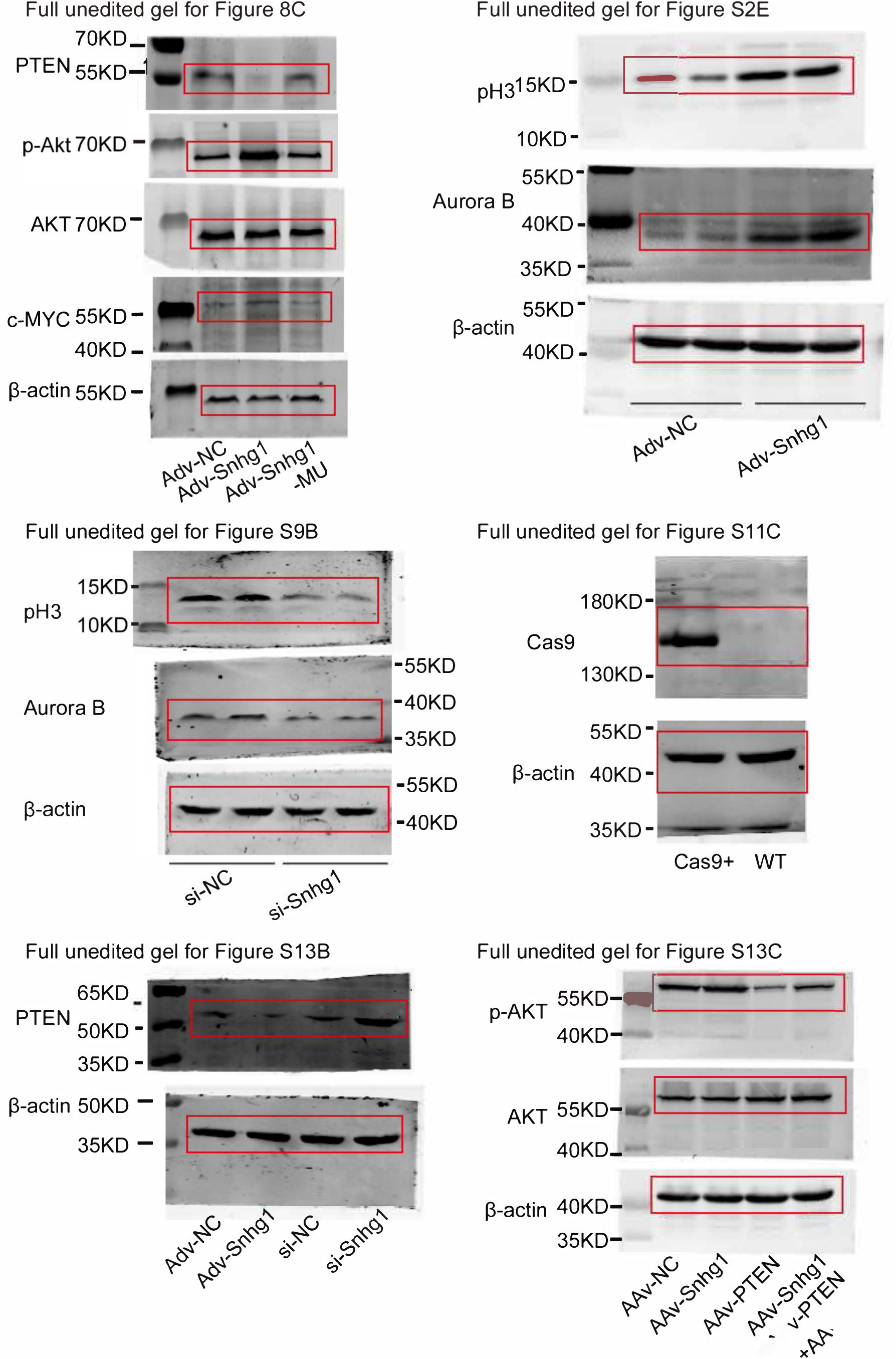
Entire unedited gel for representative cropped gels of Fig.8C,S2D,S9B,S11C,S13B,S13C. Representative cropped gels are marked by red box.

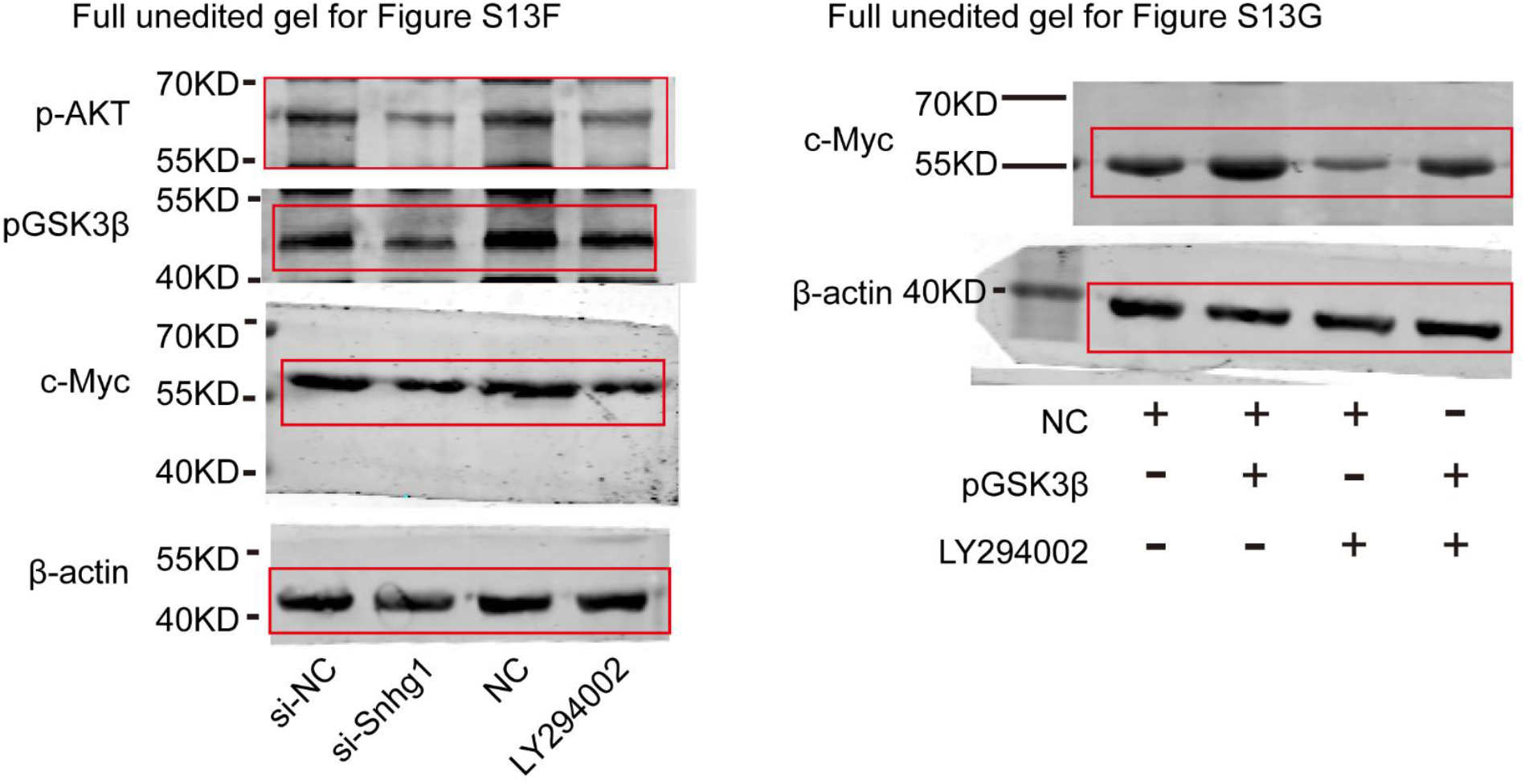
Entire unedited gel for representative cropped gels of Fig.S13F,S13G. Representative cropped gels are marked by red box.

